# Human 3D epithelioids enable continuous long-term clonal evolution studies across multiple epithelial tissues

**DOI:** 10.64898/2026.06.07.730560

**Authors:** Inês S. Ferreira, Alberto Pradilla-Dieste, Jose Antonio Valverde-Lopez, Federico Abascal, Yoshihiro Ishida, Marta Baselga, Moritz Przybilla, Andrew Lawson, Pantelis Nicola, Adrian Baez-Ortega, Mario Soriano-Navarro, Richard Butler, Michelle Miniter, Carmen Sancho-Serra, Kasandra Malasi, Chang-Bon Man, Malcolm Cameron, John A Tadross, Amy Bates, Glenn Harden, Will Ince, Gill Barnet, Rajesh Jena, Roser Vento-Tormo, Jack Martin, Krishnaa T Mahbubani, Kourosh Saeb-Parsy, Alberto J. Schuhmacher, Inigo Martincorena, David Fernandez-Antoran

## Abstract

Modeling human epithelia *in vitro* remains challenging because current systems do not fully preserve the combination of architecture, heterogeneity, clonal composition, and long-term dynamics shown *in vivo*. While 3D approaches such as organoids and organotypic cultures capture important aspects of lineage differentiation and niche signaling, they often lose stable organization over time, limiting studies of long-lasting processes such as clonal evolution and cell competition. Here, we present human epithelioids as continuous long-term, 3D epithelial cultures efficiently derived from eight adult human epithelia, including trachea, skin, buccal mucosa, esophagus, blader, urethra, submandibular gland and endometrium. Using immunostaining, electron microscopy, single-cell RNA sequencing, functional assays and somatic mutation analyses, we deeply characterized human epithelioids and confirmed that they recapitulate native architecture and cell diversity, sustain regenerative capacity, and preserve donor-specific mutational landscapes, establishing a robust and versatile platform for longitudinal interrogation of clonal evolution, tissue dynamics and responses to clinically relevant perturbations, including radiotherapy and chemotherapy, over extended timescales.

## Introduction

Human epithelia form polarized, tightly connected sheets of cells that line organs and body surfaces, providing barrier function, coordinating exchange, secretion, sensing, and defense while undergoing continuous turnover and repair^1–3^. Different epithelia show distinct architectures (e.g. stratified, pseudostratified, simple, glandular), cell type compositions, and self-renewal dynamics, typically sustained by stem or progenitor cells.

A defining feature of adult human epithelia is that they are evolving populations. Deep sequencing and clonal analyses have revealed pervasive somatic mosaicism and widespread positive selection in histologically normal tissues, frequently involving cancer-driver genes^4–7^. These studies have shown that normal epithelia contain large numbers of small mutant clones with significant heterogeneity across individuals with different exposures and cancer risk factors. Understanding how different exposures and treatments alter the clonal composition of a tissue offers new opportunities to understand early carcinogenesis and develop novel cancer prevention strategies, but this remains difficult from studies of human tissues shaped by a lifetime of complex exposures^8^.

Currently there is a lack of *in vitro* models that preserve epithelial architecture and spatial continuity while supporting stable, long-term homeostasis to study clonal evolution experimentally. Conventional 2D cultures are accessible and easy to perturb but lack tissue architecture, show altered differentiation programs and fail to replicate the mechanical conditions that influence how cells behave in the body^9^. Organoids provide a powerful access to human epithelial biology, capturing lineage specification and cell-intrinsic programs in 3D^10–13^ and can be useful to quantify mutational processes in adult stem cells^14,15^. However, their geometry limits modeling of competition across extended landscapes and their maintenance usually relies on dissociation and passaging which might reduce clonal diversity and impose culture-specific selection^16–18^

Organotypic cultures reproduce better stratification and epithelial-stromal interactions but rely on complex niche components and are generally limited to short culture times^19–22^. As a result, continuous processes that unfold over extended timescales, including somatic mutation accumulation, clonal competition or early tumor evolution remain difficult to interrogate experimentally *in vitro*^23^.

To address these limitations, we previously developed a complementary culture system termed “epithelioids”, which harnesses the intrinsic regenerative capacity of adult mouse esophageal epithelium to generate self-sustaining, 3D cultures that preserve tissue architecture and clonal dynamics over long periods without passaging^24,25^. Here, we build on this foundation to establish, optimize and characterize human epithelioids from multiple adult human tissues representing different epithelial subtypes (pseudostratified, squamous stratified, transitional, glandular and simple columnar). Human epithelioid 3D cultures provide a versatile platform that preserves epithelial architecture, displays long-term homeostasis and enables longitudinal studies of how different perturbations, including clinical therapies such as radiotherapy and chemotherapy, influence clonal dynamics and tissue toxicity in human epithelia.

## RESULTS

### Human epithelioids recapitulate the structure and cell type composition of multiple epithelia

We first aimed to determine whether long-term self-sustaining 3D epithelioid cultures could be efficiently established from a range of human tissues. Following optimization of the original mouse culture protocol, we were able to establish 3D epithelioids with high success rates from 8 human epithelia: trachea, skin, buccal mucosa, esophagus, bladder, urethra, submandibular salivary gland (SMG), and endometrium (Figure 1A). Once plated on porous transwell inserts, 1mm^2^ epithelial explants underwent cellular outgrowth, forming a new multilayered 3D epithelial structure (hereafter termed epithelioid) that covered the whole surface within two to three weeks. The epithelioids were subsequently cultured in air-liquid interface (ALI) (unless otherwise indicated) and maintained in continuous self-sustaining culture for 2, 3, 6 or 8 months (Figures 1B, 7Q, 7R and Methods). Compared to the original mouse protocol, we introduced ALI conditions and retained the original epithelial explants throughout the entire culture period to achieve a more robust long-term maintenance of human epithelioids and improve differentiation. To characterize the epithelial architecture and cell type composition in these cultures, we first performed immunofluorescence staining in 1.5- to 2-month-old epithelioids and compared them to matched donor tissues using cryosections. Epithelioids retained key architectural and molecular features of their tissue of origin, including the expression of characteristic epithelial markers, a proliferative basal layer, differentiated suprabasal compartments, and correct spatial localization of lineage-specific markers (Figures 1, 2, and S1, S2).

**Figure 1:**
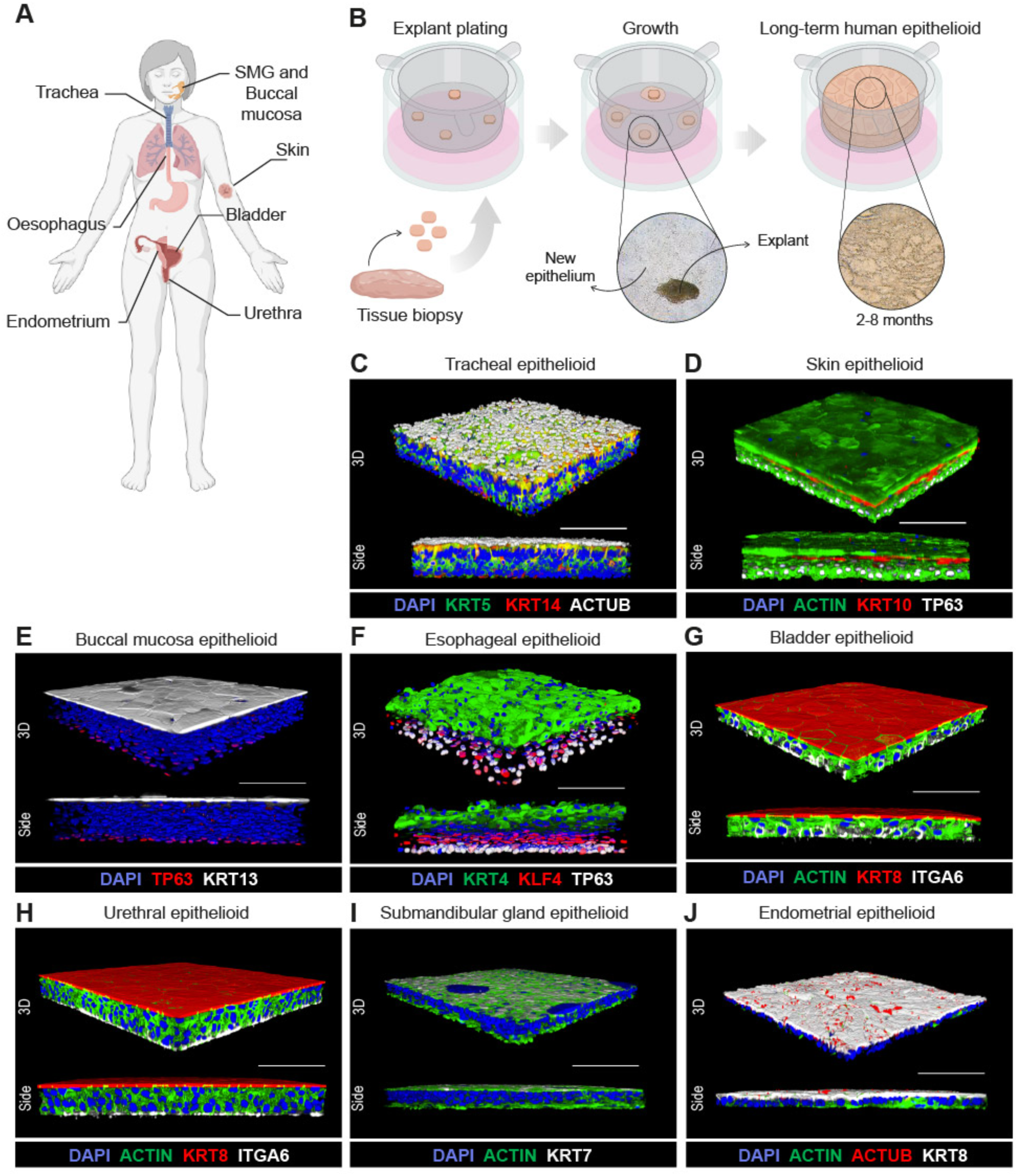
Structural and phenotypic characterization of human epithelioids generated from a diversity of epithelia. **(A)** Schematic of epithelial tissues used to generate human epithelioids. **(B)** Schematic of the setup and maintenance of human epithelioids. **(C – J)** Rendered 3D confocal images including different staining combinations to show the architecture and cell populations of human tracheal **(C)**, skin **(D)**, buccal mucosa **(E)**, esophageal **(F)**, bladder **(G)**, urethral **(H)**, submandibular gland **(I)** and endometrial **(J)** epithelioids. Scale bar, 100 μm.

**Figure 2:**
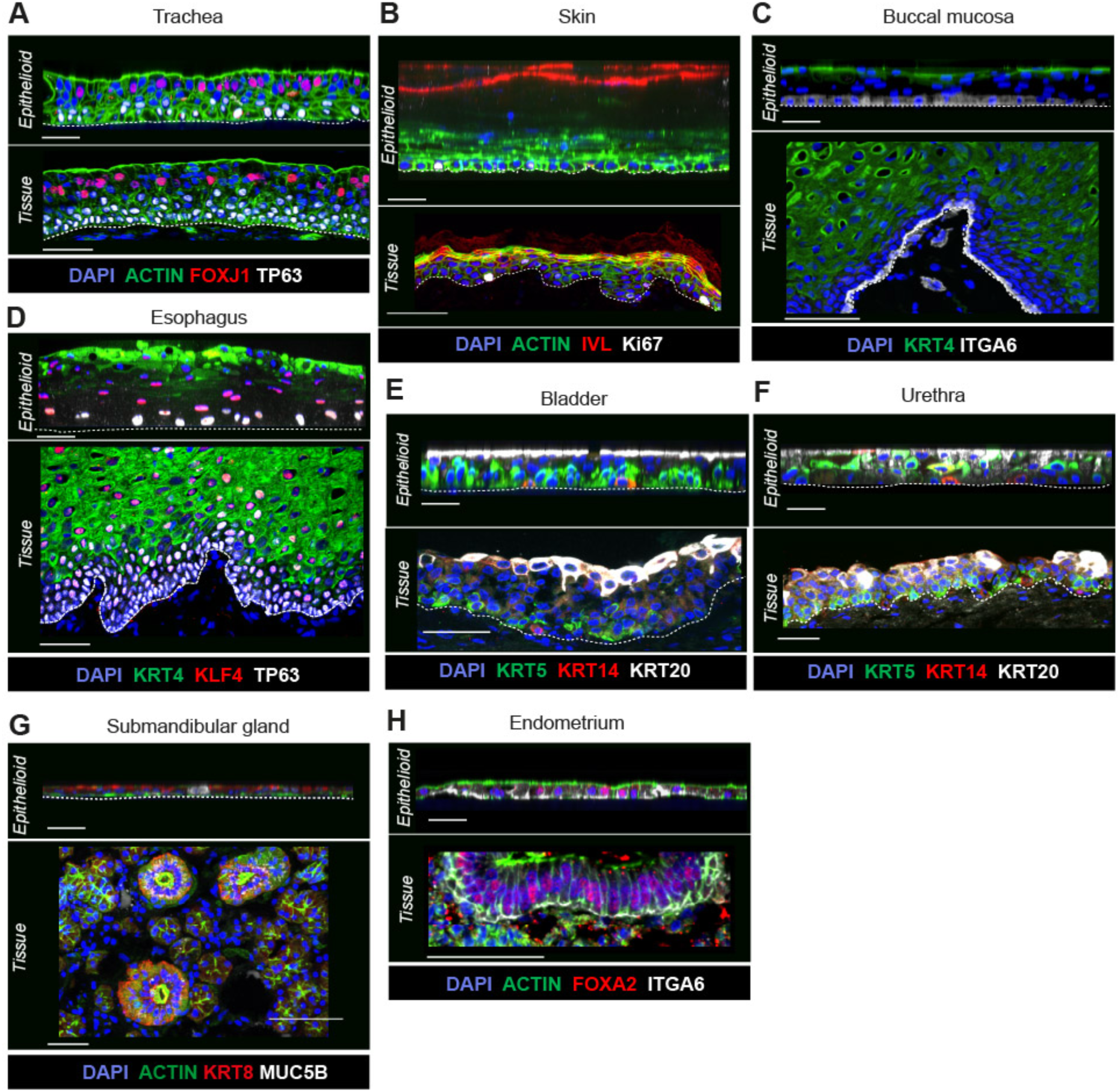
Human epithelioids recapitulate the structure and cellular distribution of human epithelia. **(A – H)** Images showing side views from 3D reconstructions of human tracheal **(A)**, skin **(B)**, buccal mucosa **(C)**, esophageal **(D)**, bladder **(E)**, urethral **(F)**, submandibular gland **(G)** and endometrial **(H)** epithelioids benchmarked against tissue cryosections. The dotted line denotes epithelioids or epithelia basal boundaries. Scale bar, 50 μm.

Tracheal epithelioids displayed a pseudostratified architecture with basal progenitors expressing KRT14, KRT5, and TP63^26,27^; ciliated cells marked by FOXJ1^28,29^ and acetylated tubulin (ACTUB)^30^; and secretory Goblet and Club cells expressing MUC5AC and SCGB1A1, respectively^31^ (Figures 1C, 2A, and S1A, S1B, S2A). Cilia organization and beating activity were comparable to native tracheal epithelium (Supplementary Video 1).

Stratified epithelioids derived from skin, buccal mucosa, and esophagus reproduced their characteristic squamous organization. Cells in the basal layer expressed progenitor markers such as TP63, ITGA6 and the proliferation marker KI67^32,33^. Skin epithelioids displayed the suprabasal markers KRT10 and IVL, characteristic of spinous and granular strata, respectively. Buccal mucosa and esophageal epithelioids showed expression of KRT13 and KRT4 in suprabasal and terminally differentiated layers. Additionally, KLF4 was found in suprabasal cells in esophageal epithelioids^34–37^ (Figures 1D-1F, 2B-2D and S1C-S1E, S2B-S2D).

Bladder and urethra epithelioids recapitulated the transitional stratified urothelium^38^, composed of basal cells expressing ITGA6 and KRT5^39^ and umbrella cells (KRT8^+^ and KRT20^+^)^40^ (Figures 1G-1H, 2E-2F, and S1F, S2E-S2F). A discrete KRT14⁺KRT5⁺ basal population corresponding to a stem cell subset found *in vivo*^41,42^ was also observed (Figures 2E and 2F).

SMG epithelioids contained ductal cells marked by KRT7 and KRT8^43^ (Figures 1I and 2G) and acinar-like cells expressing MUC5B and MIST1^43,44^ (Figures S1H and S2G), but lacked the typical glandular tubular structure found *in vivo*.

Endometrial epithelioids adopted a simple epithelium architecture with appropriate apicobasal polarity and expressing canonical epithelial and polarity markers, including KRT8, ITGA6, and E-cadherin, together with the presence of sparse ACTUB^+^ multiciliate cells^45^ (Figures 1J and S1I, S2H). We also detected cells expressing FOXA2, a transcription factor required for glandular morphogenesis *in vivo*^46,47^ (Figure 2H).

Together, these data demonstrate that human epithelioids recapitulate the structural organization, cellular composition, and marker expression profiles of their corresponding tissues of origin across different epithelial subtypes.

### Ultrastructural analysis reveals tissue-specific features and epithelial integrity in human epithelioids

To further investigate the extent to which human epithelioids resemble their tissue of origin, we evaluated their ultrastructural composition by conducting transmission electron microscopy (TEM; cross-sections) and scanning electron microscopy (SEM; surface topology) on 1.5- to 6-month-old epithelioid cultures maintained under ALI conditions. Across tissues, ultrastructural analysis revealed hallmark features of epithelial architecture, integrity and differentiation, including hemidesmosomes anchoring basal cells^48^, prominent cell interdigitations (particularly among basal cells), abundant desmosomes within intermediate layers^49^, and well-formed apical junctions sealing the luminal surface^50^. In addition, epithelioids exhibited clear epithelial polarization within differentiated suprabasal layers and contained tissue-appropriate subcellular structures and organelles, including lamellar bodies and secretory granules, consistent with terminal differentiation programs^51,52^ (Figures 3 and S3).

**Figure 3:**
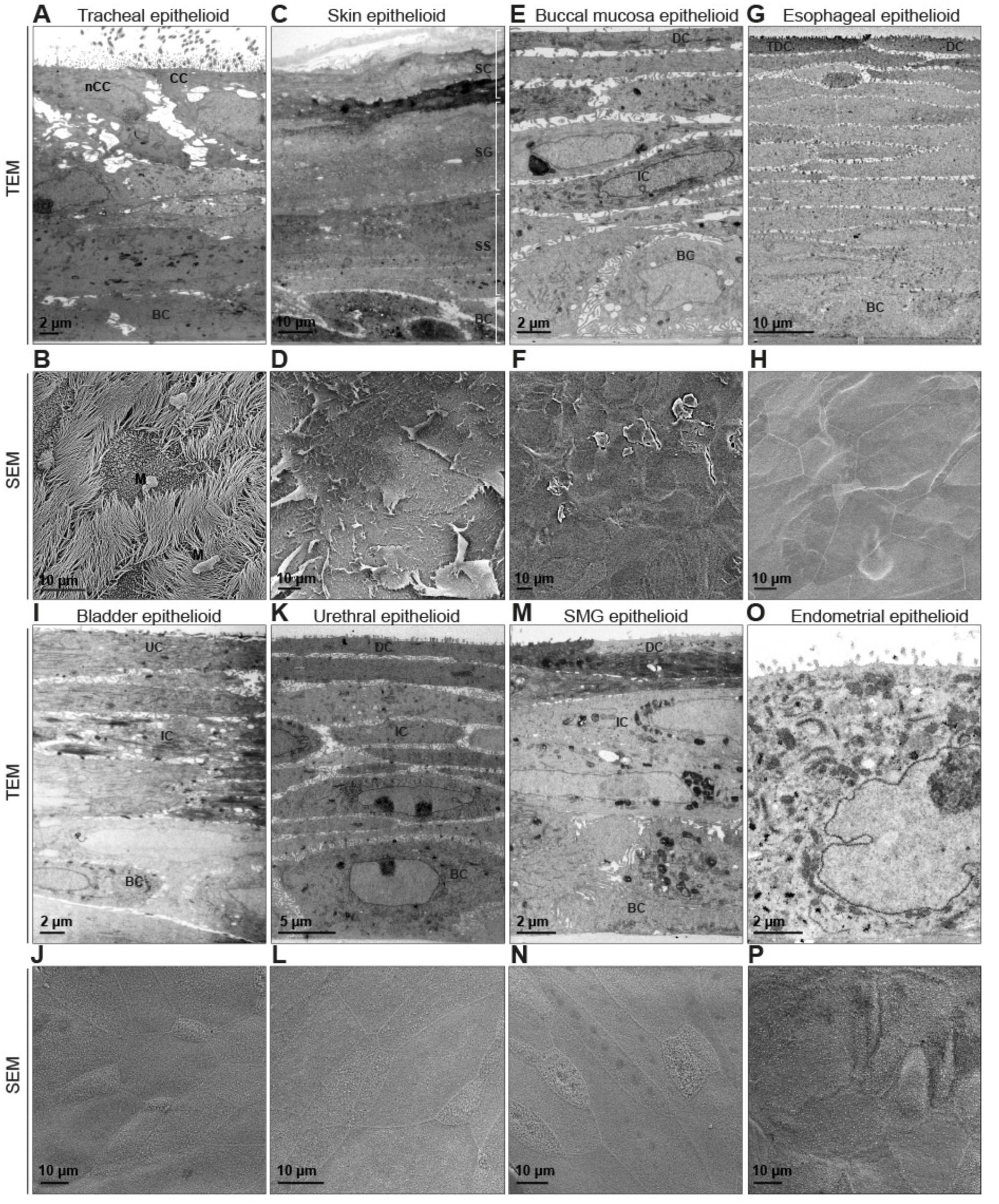
Ultrastructural analysis of human epithelioids reveals apical-to-basal organization and epithelial integrity. **(A – P)** Representative images of TEM (top) and SEM (bottom) from tracheal **(A – B)**, skin **(C – D)**, buccal mucosa **(E – F)**, esophageal **(G – H)**, bladder **(I – J)**, urethral **(K – L)**, submandibular gland **(M – N)** and endometrial **(O – P)** epithelioids (BC: basal cells; CC: ciliated cells; DC: differentiated cells; IC: intermediate cells; nCC: non-ciliated cells; SC: stratum corneum; SG: stratum granulosum; SS: stratum spinosum; UC: umbrella cells).

Pseudostratified tracheal epithelioids, showed abundant number of ciliated epithelial cells (CC) with a high density of cilia on the apical surface (Figures 3A, 3B and S3A top, S3B) with diameters of 244 nm +/-24 nm and 0.54 cilia/µm^53,54^. They also showed patterns of mitochondria accumulation beneath their surface, similar to *in vivo*^55,56^ (Figure S3A). In Goblet cells (GC), the presence of mucus could be clearly observed by electron-lucent granules (Figure S3A bottom), together with pieces of mucus being transported by cilia (Figures 3B and S3B).

Stratified squamous epithelioids derived from skin, buccal mucosa, and esophagus exhibited well-organized basal layers with prominent interdigitations and progressive suprabasal differentiation^57^ (Figures 3C, 3E, and 3G). They developed multiple layers of differentiated cells, ranging from approximately seven layers in buccal mucosa epithelioids (Figure 3E) to 12-18 layers in skin and esophageal epithelioids, respectively (Figures 3C and 3G). Keratinization was observed in skin epithelioids (Figure 3C), together with well-formed corneodesmosomes^58–60^ at cell-cell junctions (Figures S3C, top; S3E, bottom; and S3G, bottom) and flattened and densely packed fully differentiated cells at the top^61^ (Figures 3D, 3F, 3H and S3C bottom, S3D, S3F, S3H).

Urothelium epithelioids from bladder and urethra showed stratification that recapitulated the three main layers found *in vivo*: basal cells, intermediate cells and umbrella cells^62–64^ (Figures 3I, 3K and S3K top). Junction complexes with adherent junction and desmosomes (Figure S3I top) and interdigitations between cells were also clearly observed (Figures S3I bottom, S3I top and S3K), especially on urethra basal cells (Figure S3K bottom).

Submandibular salivary gland epithelioids (SMG) exhibited densely packed intermediated cells with numerous interdigitations but less evident desmosomes^65,66^ (Figure 3M). Secretory granules were frequently found (S3M top). Cell-cell interactions and polarization were preserved across samples, indicating the ability of SMG epithelioids to partially restore native epithelial organization *in vitro* (Figure S3M bottom).

Endometrium epithelioids displayed a simple epithelial architecture with a clear apicobasal polarity. The apical surface exhibited cytoplasmic protrusions consistent with those described in native endometrial luminal epithelium^67,68^ (Figures 3O, 3P and S3O bottom) and the presence of sparse ciliated cells (Figure S3P). Secretory granules were also observed in the cytoplasm (Figure S3O).

Altogether these ultrastructural analyses demonstrate that human epithelioids preserve key architectural and subcellular features of epithelial tissues, required for proper epithelial integrity, polarity and tissue function.

### Single-cell sequencing reveals that human epithelioids recapitulate the epithelial diversity of tissues

Human epithelial tissues are characterized by cellular heterogeneity. Single-cell sequencing has been used to describe the prominent cell types of human tissues, driven by large-scale initiatives such as the Human Cell Atlas (HCA)^69^. To evaluate the presence of the main epithelial cell types in human epithelioids, we performed single-cell RNA sequencing^70–72^ (scRNA-seq) on 1.5- to 2-month-old epithelioid cultures derived from six distinct human tissues: trachea, skin, buccal mucosa, esophagus, bladder, and submandibular salivary gland (SMG) (Figure 4 and Figure S4). Following mild enzymatic dissociation and microfluidic capture, we retained a total of 41,494 high-quality single-cell transcriptomes for downstream analysis. We leveraged literature-derived marker genes to annotate all epithelioid cultures. To complement this approach, we integrated published cell type gene expression signatures from reference datasets (Figures 4A, 4E, 4I, 4M, 4Q and 4U), method^73^, trachea^74^, bladder^75^, skin^76^, oral mucosa^77^, esophagus^78^, submandibular gland^79^.

**Figure 4:**
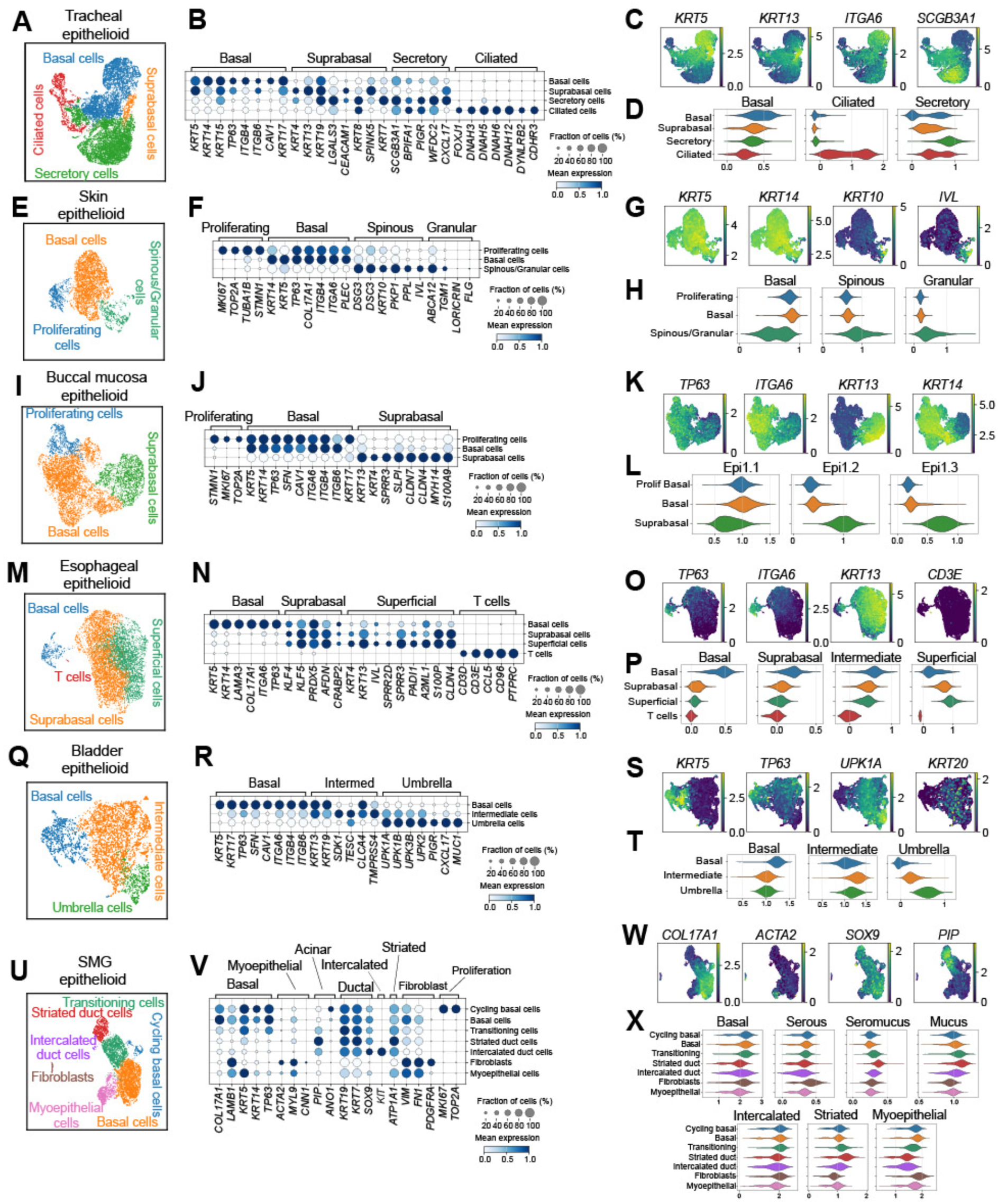
Single-cell transcriptomics confirm the protein-level data and expose tissue-resident immune cells in human epithelioids. **(A, E, I, M, Q, U)**, Uniform manifold approximation and projection (UMAP) plots of scRNA-seq data from **(A)** tracheal (n=12,423), **(E)** skin (n=4,401), **(I)** buccal mucosa (n=7,353), **(M)** esophageal (n=7,528), **(Q)** bladder (n=3,897), **(U)** submandibular gland (n=5,892) epithelioids. **(B, F, J, N, R, V)** Dot plots illustrating the scaled, mean expression and detection frequency of selected markers genes across all clusters identified from single-cell RNA sequencing analyses in human tracheal **(B)**, skin **(F)**, buccal mucosa **(J)**, esophageal **(N)**, bladder **(R)** and submandibular gland **(V)** epithelioids. **(C, G, K, O, S, W)** UMAP plots showing normalized expression (counts per 10,000 reads; CP10K) of selected marker genes in tracheal **(B)**, skin **(E)**, buccal mucosa **(H)**, esophageal **(K)**, bladder **(N)** and submandibular gland **(Q)** epithelioids. **(D, H, L, P, T, X)** Violin plots showing gene score expression patterns across cell clusters, calculated from gene sets identified in previously published studies.

Across all tissues, epithelioids recapitulated the major cell populations expected from the original epithelia. Tracheal epithelioids (Figures 4A-4D and S4G, S4H) comprised four principal epithelial lineages defined by canonical markers: ciliated (*FOXJ1*, *CDHR3)*, suprabasal (*KRT8*, *KRT19*), secretory (*SCGB3A1*, *WFDC2*), and basal (*KRT5*, *KRT14*, *ITGA6*, *MKI67*).

Skin epithelioids (Figures 4E-4H and S4I, S4J) displayed three principal keratinocyte populations corresponding to basal progenitors (*KRT14*, *KRT5*, *ITGA6*), a proliferative basal compartment (*KRT14*, *KRT5*, *MKI67*, *TOP2A*), and differentiated suprabasal keratinocytes representing the spinous/granular layers (*DSG3*, *IVL*). Consistently with the anucleate state of terminally differentiated corneocytes (shown in Figures 2B and S1C), cells corresponding to the cornified layer were not detected in this dataset^80^.

Buccal mucosa epithelioids (Figures 4I-4L and S4K, S4L), delineated three well-defined epithelial populations corresponding to basal progenitors (*TP63*, *ITGA6*, *KRT14*), a proliferative basal compartment (*TP63*, *ITGA6, KRT14*, *MKI67*), and a differentiated suprabasal keratinocytes characteristic of the buccal mucosal epithelium (*KRT13*, *KRT4*, *SPRR3*).

Similarly, esophageal epithelioids (Figures 4M-4P and S4M, S4N) displayed basal (*TP63*, *ITGA6, KRT5, COL17A1*), suprabasal (*KLF4*), and superficial (*SPRR3*, *KRT4*, *KRT13, IVL*) populations, with a subset of proliferating basal cells expressing *MKI6*7 (Figure S4M)^81–84^.

Bladder epithelioids (Figures 4Q-4T and S4O, S4P) reproduced the main urothelial compartments of the human urothelium, including basal (*KRT5*, *ITGA6*, *TP63*), intermediate (*KRT13*, *KRT19*, *SDK1*), and umbrella (*UPK1A*, *UPK2*) cell layers^40^.

SMG epithelioids exhibited the highest degree of cellular heterogeneity (Figures 4U-4X and S4Q, S4R), encompassing basal (*COL17A1*, *KRT14*, *TP63*), proliferating basal (*COL17A1*, *KRT14*, *TP63*, *MKI67*), myoepithelial (*MYL9*, *ACTA2*), intercalated ductal (*KIT*, *SOX9*, *KRT7*, *KRT19*), striated ductal (*ATP1A1*, *PIP*) together with a fibroblast-like population expressing mesenchymal markers (VIM, FN1, PDGFRA). Although acinar cells were detected by immunofluorescence (Figures 1SH and 2G), this population was not resolved in the scRNA-seq dataset, consistent with their low abundance in the culture and perhaps technical limitations in capturing highly secretory cell states^85^.

Remarkably, we identified a discrete cluster of T cells (*CD3D*, *CD3E*, *CCL5*) in esophageal epithelioids (Figure 4M and 4N). To rule out possible residual RNA contamination from the original explants, we applied in silico correction approaches (Methods). The T cell signal remained prevalent and clearly abundant in our data. We performed complementary IF staining for CD45 and CD3E, confirming the presence of T cells within the suprabasal compartment of the human epithelioid culture (Figure S4A).

These data indicate that immune cells infiltrate esophageal epithelioids cultures and remain viable for at least 1.5 to 2 months in continuous culture. We also extended our IF analysis to other epithelioid samples and found comparable immune cell infiltration in tracheal, buccal mucosa, bladder, urethral and SMG epithelioids after 1.5 to 2 months in culture (Figures S4B-S4F). While this observation will require further characterization, our results highlight that human epithelioids preserve the major epithelial cell types and aspects of the native tissue immune landscape.

### Human epithelioids maintain tissue-like organization and homeostatic dynamics over time

Proliferative activity and cellular differentiation are key features of epithelial turnover. A common approach to quantify this is 5-ethynyl-2′-deoxyuridine (EdU) labeling^24,86^. We performed EdU labeling on 1.5 to 2-month-old epithelioids cultured under ALI conditions, with EdU incorporation being measured at 1 hour or 48 to 96 hours post-incubation to assess proliferation and differentiation dynamics within each human epithelioid culture, respectively (Figures 5 and S5).

**Figure 5:**
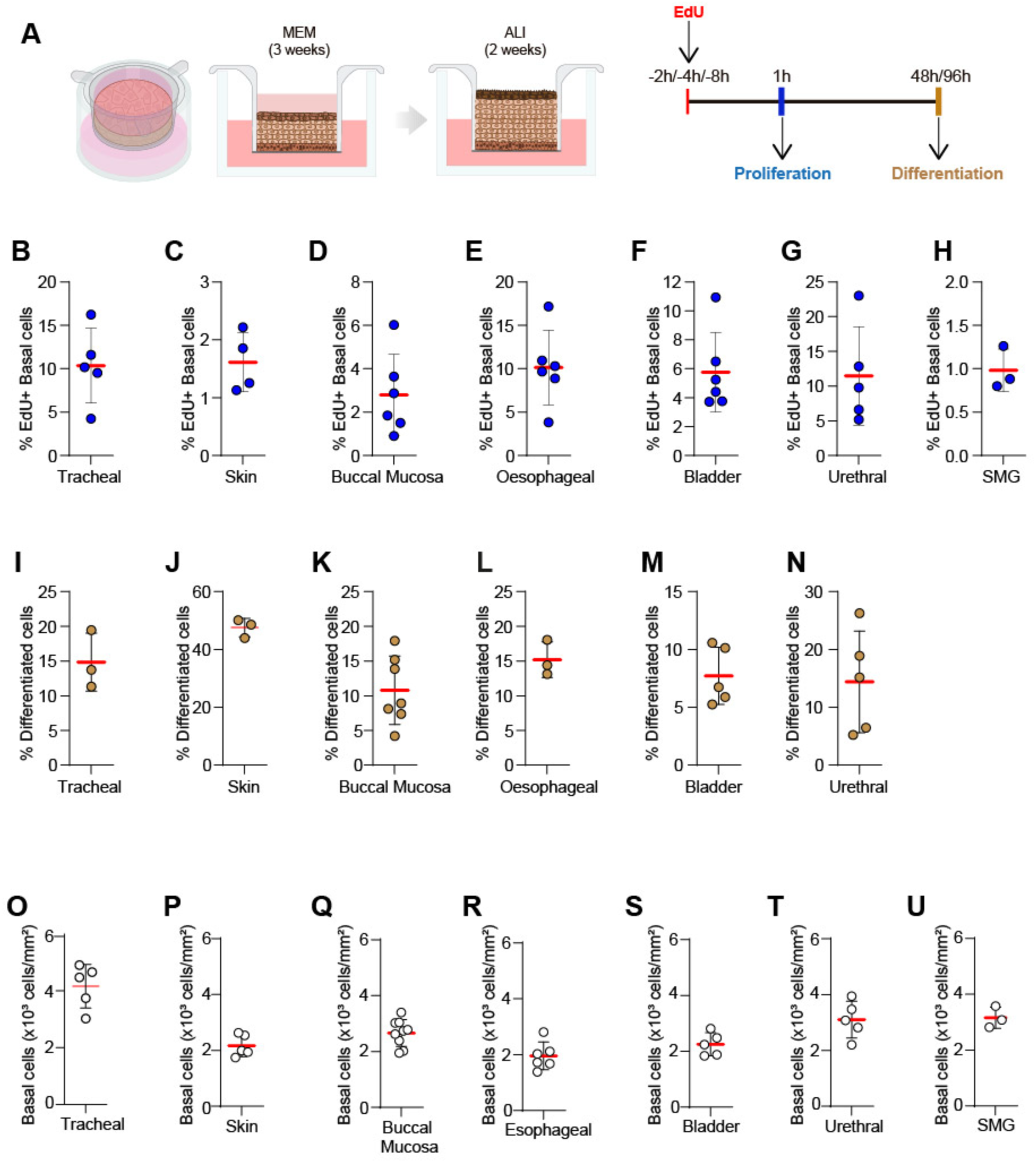
Human epithelioids recapitulate native tissue dynamics of cell proliferation and differentiation. **(A)** Schematic of the EdU labeling assay used to measure proliferation, differentiation, and basal cell density in human epithelioids. **(B – I)** Percentage of basal EdU^+^ (proliferating) cells per total number of basal cells. **(J – O)** Percentage of suprabasal EdU^+^ (differentiated) cells relative to total EdU^+^ cells. **(P – W)** Basal cell density expressed as total number of basal cells per mm^2^. Results are expressed as mean plus SD from at least three different donors.

To evaluate proliferative capacity, epithelioids were incubated with EdU for 2, 4 or 8 hours, and incorporation was analyzed by confocal microscopy one hour later. Across all epithelioid types, EdU labeling was confined to the basal compartment, consistent with the localization of progenitor- or stem-like cells shown in Figures 1 and 2. Quantification of EdU⁺ basal cells revealed distinct proliferative profiles across tissues. Submandibular salivary gland, skin, and buccal mucosa epithelioids showed the lowest mean frequencies of EdU^+^ basal cells, at 0.98, 1.61, and 2.31%, respectively (Figures 5C, 5D, 5H and, S5C, S5D, S5H). Bladder epithelioids displayed an intermediate level of proliferation, with an average of 5.76% EdU^+^ basal cells (Figures 5F and S5F). By contrast, urethral, tracheal, and esophageal epithelioids were the most proliferative, with 11.49, 10.36, and 10.13% EdU^+^ basal cells, respectively (Figures 5B, 5E, 5G and S5B, S5E, S5G). Notably, the proliferative indices measured in epithelioids fell within those ranges previously reported for the corresponding human tissues ^87–89^.

Next, we investigated the differentiation capacity across stratified epithelioid cultures. To do so, we conducted EdU pulse-chase experiments. An EdU pulse was administered for 2, 4 or 8 hours and labelled suprabasal cells were quantified 48 or 96 hours later (Figure S5A).

Tracheal, buccal mucosa, esophageal, and urethral epithelioids showed relatively low differentiation indices, with an average of approximately 10–16% of total EdU^+^ suprabasal cels (Figures 5J, 5L, 5M, 5O and S5J, S5L, S5M, S5O). Bladder epithelioids showed a lower mean differentiation index of 7.11% (Figures 5N and S5N). In contrast, skin epithelioids displayed a substantially higher proportion of suprabasal EdU^+^ cells, with an average of 47.56% (Figures 5K and S5K).

Finally, we quantified the basal cell density across human epithelioids and observed marked tissue-specific variation, with densities ranging from approximately 4000 basal cells per mm² in tracheal epithelioids (Figures 5P and S5P), 3000 basal cells per mm² in urethral and SMG epithelioids (Figures 5U, 5V and S5U, S5V), to 2000 basal cells per mm^2^ in skin, buccal mucosa, esophageal and bladder epithelioids (Figures 5Q-5T and S5Q-S5T).

Together, these results demonstrate that human epithelioids retain key proliferative and differentiation features of their native epithelia, and might support their use as long-term, tissue-like *in vitro* system for studying epithelial homeostasis.

### Human epithelioids maintain barrier homeostasis and restore integrity after injury

An essential hallmark of epithelial tissues is their ability to establish a selective barrier and to rapidly restore tissue integrity following injury^90,91^. To evaluate whether human epithelioids recapitulate these core physiological properties, we evaluated barrier function and wound-healing capacity in 2-month-old cultures (Figures 6 and S6).

**Figure 6:**
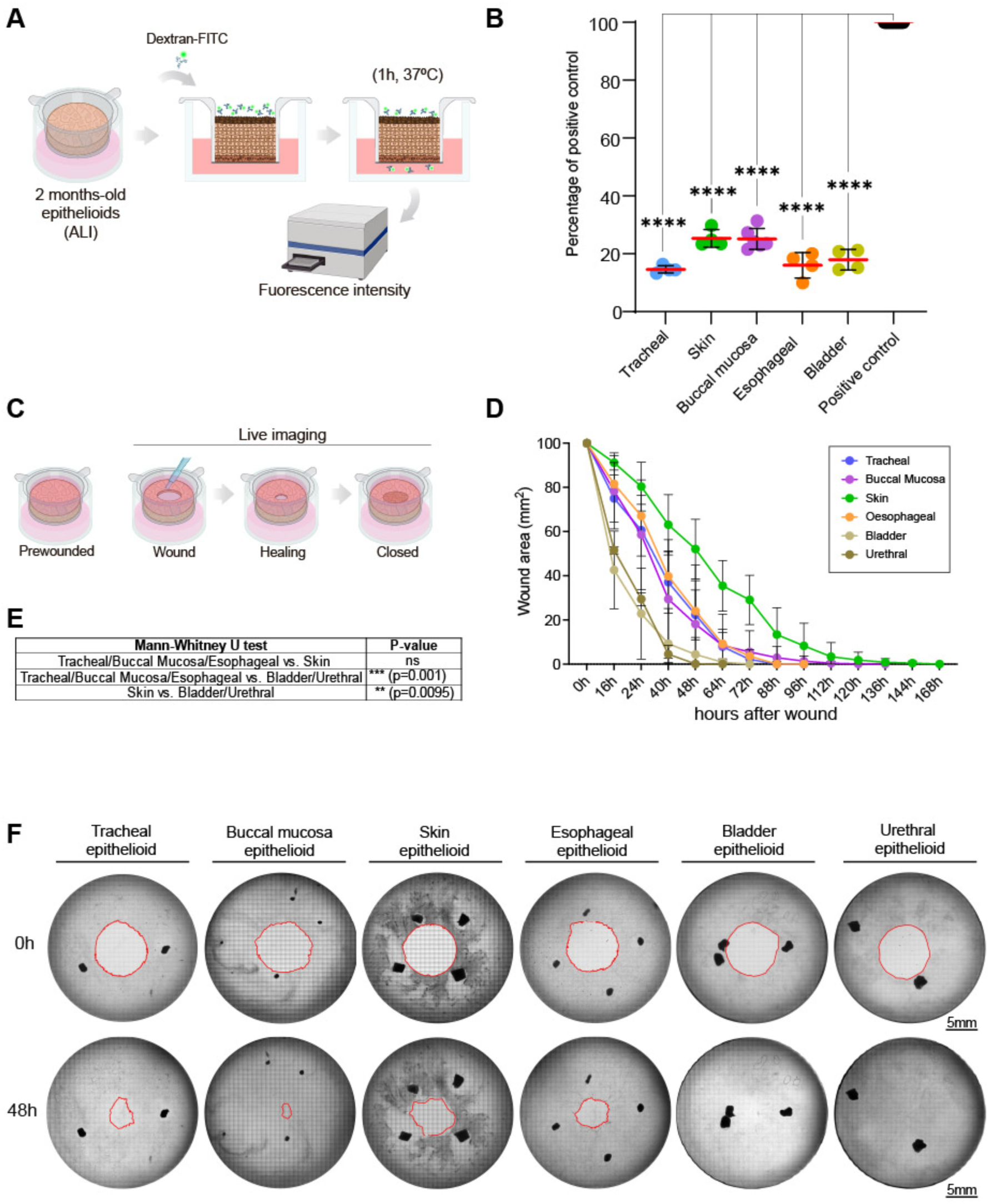
Human epithelioids maintain barrier homeostasis and wound healing capacity after injury. **(A)** Schematic of the Dextran-FITC permeability assay used to assess barrier integrity in human epithelioids. **(B)** Fluorescence intensity in human epithelioids expressed as a percentage of paired positive controls. Results are expressed as mean plus SD from at least three different donors. **(C)** Schematic of the live imaging setup used to assess healing capacity after injury in human epithelioids. **(D)** Wound area in human epithelioids over time, expressed as a percentage of the initial wound area for each epithelioid. Results are expressed as mean plus SD from at least three different donors. **(E)** Comparison of healing rates between different human epithelioids. Kruskal-Wallis test was applied for statistical comparison of the wound closure rates among the three groups; (**, p=0.001; ***, p=0.0006). **(F)** Brightfield images of human epithelioids immediately after wounding (0 h) and 48 h post-wounding. Red lines outline wound areas.

Barrier integrity was measured using paracellular permeability with a 4 kDa fluorescein (FITC)-dextran flux assay, a widely used *in vitro* readout of tight-junction function and overall epithelial barrier performance^92–99^. This approach is particularly sensitive to paracellular leakage and provides a quantitative measure of functional barrier integrity.^100^

4 kDa FITC-dextran was added to the apical compartment of ALI epithelioid cultures derived from human trachea, skin, buccal mucosa, esophagus, and bladder, and permeability was assessed after one hour (Figure 6A). All epithelioid cultures exhibited considerably reduced FITC-dextran passage relative to empty-well controls (Figure 6B). Across the five epithelioid types analyzed, average permeability was reduced to 19.1% of the empty-well control (range, 17-21%), demonstrating that human epithelioids recapitulate a functional barrier.

Next, we investigated whether epithelioids engage in a wound-repair response. We implemented a standardized punch-wound assay and quantified wound closure dynamics over time^33,101–104^. Epithelioids generated from trachea, buccal mucosa, skin, esophagus, bladder, and urethra were wounded using a sterile 8-mm biopsy punch, and repair was tracked by transmitted-light live imaging across the entire insert (452.4 mm² tile scan) (Figure 6C).

All epithelioid types re-epithelialized the wound area and achieved complete closure within 36-168 hours, with kinetics strongly dependent on tissue of origin (Figure 6D-6F). Quantification of wound area over time revealed a reproducible hierarchy of repair rates: bladder and urethral epithelioids closed most rapidly (36-48 hours), buccal mucosa, tracheal, and esophageal epithelioids showed intermediate kinetics (72-96 hours), and skin epithelioids exhibited the slowest closure (156-168 hours) (Figure 6D-6F). Closure trajectories were consistent across biological replicates, supporting the robustness of this platform for comparative interrogation of human epithelial repair mechanisms *in vitro*^105,106^.

Proliferation contributes to epithelial wound repair across different tissues^107–112^. To evaluate the involvement of proliferation in the wound healing response of epithelioids, we quantified S-phase entry by EdU incorporation at three defined stages: pre-wounding, 16 hours post-wounding, and 7 days after wound closure (Figure S6A). Despite donor-to-donor variability, buccal mucosa, skin, esophagus, bladder, and urethra epithelioids exhibited a clear increase in the fraction of EdU⁺ cells at 16 hours (expressed as fold change relative to pre-wound), consistent with a transient proliferative response during early repair^113^ (Figure S6D-M). EdU incorporation returned to baseline 7 days after closure in these epithelioids, indicating restoration of homeostatic proliferative levels. In contrast, tracheal epithelioids displayed a distinct temporal profile, with increased EdU⁺ cells evident at 7 days after closure rather than at 16 hours (Figure S6B, S6C), pointing to tissue-specific link between wound repair and cell dynamics.

### Human epithelioids maintain the polyclonality of human epithelia

DNA sequencing studies in the last decade have revealed that as we age normal epithelia are colonized by large numbers of small mutant clones carrying somatic mutations that confer proliferative advantage (frequently mutations in cancer-driver genes). These studies have provided information on the rate of somatic mutations, the dominant mutational processes (mutational signatures), and the frequency and sizes of clones bearing positively selected driver mutations in each tissue^4–6,114–119^. The utility of mouse esophageal epithelioids for studying clonal competition has been previously demonstrated^24,25^, however, the extent to which human epithelioids recapitulate the physiological mutational landscape across human tissues remains unstudied.

We performed in-depth somatic mutation analyses of 104 cultures from 9 donors, including skin (6 epithelioids/1 donor), buccal mucosa (24/4), esophageal (26/5), tracheal (24/4), and bladder (24/4) human epithelioids (Figure 7A). To quantify mutation rates, mutation signatures and clonal landscapes in each culture, we used a high-fidelity duplex sequencing protocol (NanoSeq), which accurately detects somatic mutations in single molecules of DNA, enabling sensitive somatic mutation studies on any sample independently of clonality (Methods). Specifically, we used targeted NanoSeq with a gene panel covering the most frequent driver genes in normal human epithelia (239 genes)^119,120^. Mutation rates, signatures, and driver landscapes were compared to matched original tissue samples (n = 7) and to published studies of human tissues. In total, 237 NanoSeq libraries were sequenced to a median duplex coverage of 257 dx per library (range = 31-698 dx).

**Figure 7:**
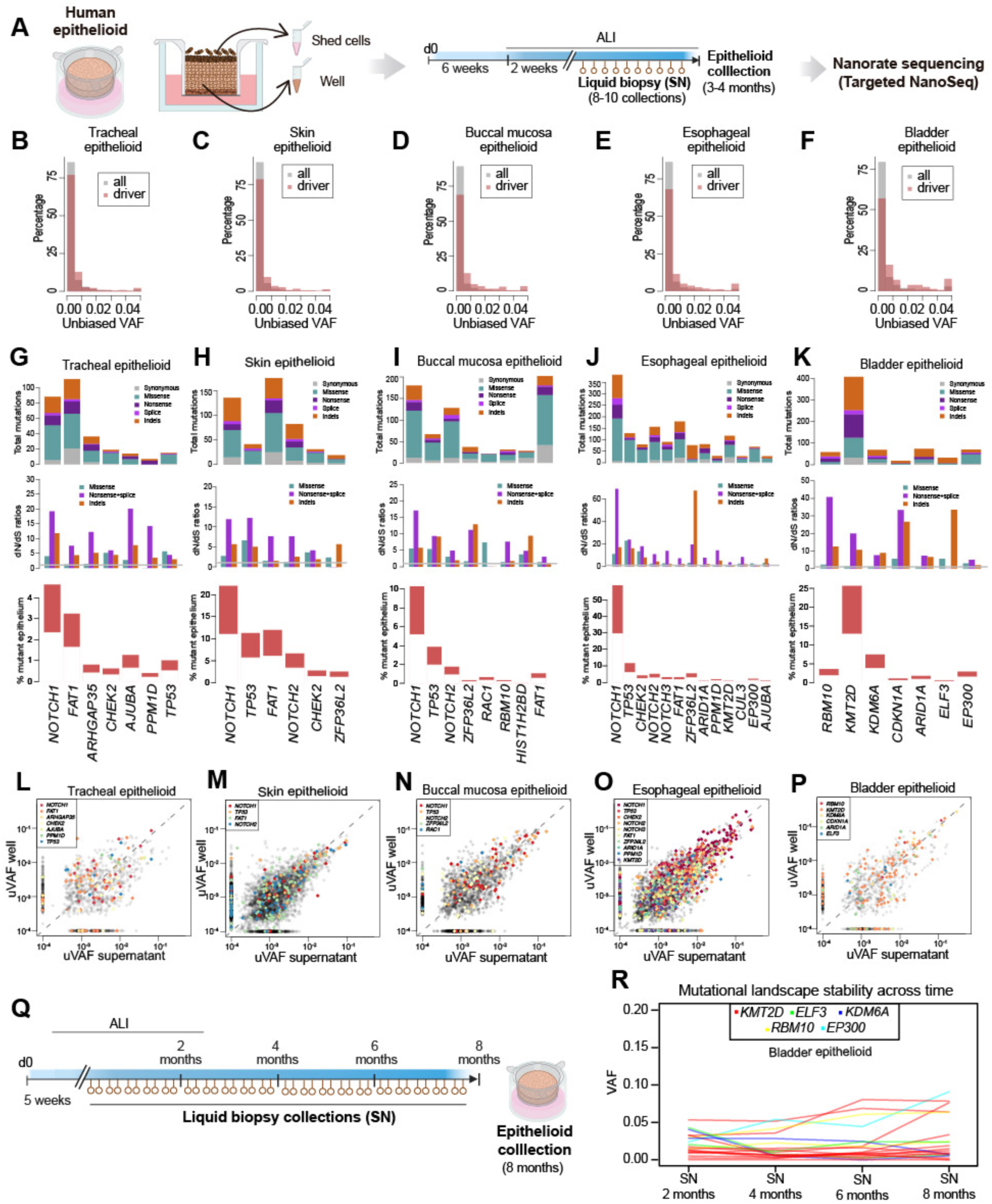
Human epithelioids recapitulates the mutational landscape of human epithelia. **(A)** Schematic of human epithelioid and supernatant collection for targeted Nanoseq. **(B-F)** Unbiased variant allele fraction (uVAF) of the mutations called with NanoSeq across epithelioids derived from five tissue types, highlighting non-synonymous mutations on driver genes in pale red. (**G-K**) Analysis of genes under positive selection in each tissue-type, showing total number and type of mutations (top), coefficients of selection or dN/dS ratios (middle) and percentage of mutant epithelium (bottom). (**L-P**) Comparison of uVAF for each mutation in the supernatant and the bulk epithelioid culture, across each of the five tissue types. (**Q**) Schematic of 8-month supernatant continuous collection of human bladder epithelioid culture (**R**) Evolution of clone sizes through time by looking at the top 20 highest VAF driver mutations.

The NanoSeq data confirmed that all epithelioid cultures were polyclonal, retaining a characteristic feature of human tissues *in vivo*. Most of the somatic mutations detected in the cultures had unbiased variant allele fractions (VAFs) below 0.5%, confirming that most cultures were composed of many small clones (Figures 7B-7F). Exceptionally, we also observed some cultures with large clones bearing known driver mutations, including one esophageal culture from a 45-year-old donor with a *NOTCH1*-mutant clone reaching a VAF of 30% (estimated to account for at least 30% of the cells in the culture) (Figure 7F). Large *NOTCH1* clones are common in normal esophagus^6^, and it is likely that this large clone dominated the original tissue explant used to establish the culture. However, it is also possible that this clone may have preferentially expanded during the generation of the culture. To explore more formally whether the generation or long-term maintenance of epithelioid cultures favors the expansion of clones carrying certain driver mutations, we next investigated the frequency of driver mutations in cultures from different tissues.

### Human epithelioids recapitulate the driver landscape of normal epithelia

Each human epithelia displays a distinct landscape of cancer-driver mutations, with substantial variation across tissues^6,118,119,121^. To study the driver landscape across epithelioids, we applied *dNdScv* to the targeted NanoSeq data from the five epithelioid types above. The resulting lists of genes under positive selection in each tissue-derived human epithelioid culture closely resembled those observed *in vivo*. *NOTCH1* emerged as the top positively selected gene in skin, esophageal, buccal, and tracheal epithelioids, consistent with previous reports performed in the corresponding tissues^6,7,118,121,122^ (Figures 7G-7J). Other recurrently selected genes included *TP53*, *FAT1*, *NOTCH2*, and *CHEK2*, each implicated in epithelial homeostasis and clonal fitness^119^. The driver landscape in bladder epithelioids included *RBM10*, *KMT2D*, *KDM6A*, *CDKN1A*, *ARID1A*, *ELF3*, *EP300*, in agreement with human sequencing studies^123,124^ (Figure 7K).

Despite occasional expansions of mutant clones in some cultures, the mean fractions of cells estimated to carry driver mutations in the cultures were similar to those observed *in vivo*, suggesting that the culture process does not greatly alter clonal selection, at least in the timescales used in our experiments. For instance, our five esophagus donors ranged between 44 and 79 years old and showed mutant cell fractions for *NOTCH1* and *TP53* ranging 11-57% and 1.6-11.8%, respectively (lower bound estimates, Methods). These figures appear broadly consistent with sequencing studies of human esophageal epithelium^6,122^. Taken together, these results show that the driver composition of epithelioids resembles that of the original epithelia, reinforcing their utility as a model for clonal competition studies in human epithelial tissues.

### Mutation rates and signatures in human epithelioids reflect *in vivo* exposures

Cell culture systems such as induced Pluripotent Stem Cells (iPSCs), organoids or single-cell-derived colonies have previously been shown to accumulate high rates of mutations *in vitro*, particularly due to oxidative damage, characterized by a high rate of G>T transversions^7,14,125,126^. This revealed that these cell culture systems under normal oxygen concentrations impose considerable oxidative stress on cells, which complicates mutagenesis studies *in vitro*^125^. In stark contrast, analysis of the mutational spectra in all the human epithelioids sequenced revealed no clear evidence of an oxidative damage signature and showed spectra consistent with those reported *in vivo*. Remarkably, this was the case even after 8 months of continuous culture (Figure S7D).

To determine how distinct mutational processes contribute to the somatic mutation burden observed in human epithelioid cultures, we then performed mutational signature analyses on the epithelioid NanoSeq data. Signature decomposition identified several dominant mutational processes largely consistent with the known mutational processes operative in the original tissue samples (Figure S7A). This mostly reflected pre-existing somatic mutations in the cells that seeded each culture. Signature A (sigA) resembled COSMIC SBS5 (cosine similarity = 0.91), a clock-like mutational signature observed across normal tissues. Signature B (sigB) corresponded to COSMIC SBS7, a signature due to UV exposure, which dominated the mutation landscape of skin epithelioids (cosine similarity = 0.95). Signature C (sigC) was highly similar to COSMIC SBS16 (cosine similarity = 0.98) (Figure. S7A), a mutational signature associated with alcohol consumption^122^. Reassuringly, SigC/SBS16 was prevalent in esophageal and tracheal epithelioid cultures (Figures S7B and S7C).

Apart from the physiological and expected mutational signatures described above, we also detected a distinctive APOBEC-like culture-specific mutational process in a minority of epithelioid cultures, Signature D (Figure S7A). This signature was characterized by TC[AT]>T substitutions, similar to COSMIC SBS2 (cosine similarity = 0.98), but not co-occurring with SBS13. This signature appeared to be more prevalent in supernatants (Figure S7B), which may suggest APOBEC activation during or after differentiation within the epithelioids. Signature E did not clearly match any single COSMIC signature and appears to be a combination of other signatures, with some contribution of SBS13 (APOBEC).

Overall, apart from the *in vitro* activation of SBS2 in a minority of cultures, the mutational profiles of the epithelioids matched those expected for their original tissues, without evidence of the high rates of oxidative damage reported in some other culture systems^127,128^. This suggests that epithelioids may be a valuable *in vitro* model for the study of endogenous mutational processes, which have traditionally been hard to study in classical culture systems.

### Non-invasive longitudinal tracking of clonal evolution using supernatants

Studying clonal competition *in vitro* could help screen and develop novel molecular prevention interventions that suppress more tumorigenic mutant clones. Human epithelioids could offer a model system which closely resembles *in vivo* dynamics of human tissues, thereby providing a valuable tool to study the impact of experimental interventions on different mutant clones. Standard collection methods require destruction of the culture, preventing longitudinal studies within a culture. Since human epithelioids continuously shed differentiated cells into the culture medium, we hypothesized that culture supernatants could provide a non-invasive readout of the clonal composition of the culture, similar to an *in vivo* liquid biopsy collection, to enable longitudinal studies.

To determine whether the clonal composition of the culture can be measured non-invasively, we performed targeted NanoSeq of supernatants and their matched cultures. Specifically, to obtain enough supernatant material to perform deep targeted NanoSeq, supernatants from 2 to 2.5 months-old epithelioids (n=92) were collected twice a week over 4 to 5 weeks. Matched cultures were harvested at the experimental endpoint (3 - 4 months; Figure 7A). Comparing the allele frequencies of all mutations detected in the wells or supernatants revealed a good concordance, supporting the validity of supernatants as a non-invasive readout of the clonal composition of a culture (Figures 7L-7P). Given the presence of dozens of detectable clones per well, this method enables parallel, longitudinal tracking of clonal composition over time.

We took advantage of the ability to sample cultures longitudinally using supernatants to track changes in mutation rates, signatures, and clonal composition during an 8-month long continuous culture of a bladder epithelioid (4 replicate cultures, each with 4 supernatant time points). This revealed remarkably stable burdens, signatures, and clonal composition despite the long time in culture (Figures 7Q, 7R and S7D, S7E). The average increase in mutation burden over 8 months in continuous culture was small (∼3.4×10^-8^ substitutions/bp, CI95%: 1.1 - 5.8×10^-8^, *P* = 0.006), with a stable mutational spectrum throughout (Figures S7D and S7E), and without major changes in the polyclonal composition (Figures 7Q and 7R).

In summary, our analyses showed that human epithelioids preserve features of somatic evolution observed *in vivo*, including polyclonal architecture, mutation burdens, and driver gene composition. We further introduce a non-invasive approach to resolve clonal composition longitudinally applying targeted NanoSeq to culture supernatants, enabling repeated, quantitative tracking of clonal dynamics without perturbing the system.

## Discussion

Our study positions human epithelioids as a long-term experimental platform for studying somatic evolution in human epithelia. This system preserves key features of tissue architecture, clonal heterogeneity, and epithelial function while enabling longitudinal sampling of clonal composition, addressing a central limitation in the field of epithelial tissue modelling and somatic evolution. Although normal epithelia are increasingly understood as dynamic evolutionary systems shaped by spatial constraints, turnover, and context-dependent selection^8,129–133^, most available approaches remain either cross-sectional or experimentally distorted by dissociation and repeated passaging^11,13,17,22,134,135^. Notably, organoid technologies, historically dominated by closed cyst-like architectures, are also beginning to shift toward more accessible open configurations^136,137^ (e.g., epithelial sheet- or layer-like formats) to improve experimental access and better reproduce key aspects of tissue biology, but still depend on dissociation and passaging. In this context, human epithelioids extend biological fidelity into a stable, continuous long-term 3D culture system with an experimentally accessible surface and compartmental organization, providing a controllable framework to connect epithelial physiology to clonal dynamics over time without repeated disruptive culture manipulation.

We have shown molecular, structural, and functional evidence supporting tissue fidelity in human epithelioids. Immunofluorescence and ultrastructure analyses showed that human epithelioids retain the architecture of their tissue of origin. Immunofluorescence and scRNA sequencing confirmed that epithelioids recapitulate the 3D organization and cellular composition of the original epithelia, while ultrastructural analyses revealed canonical epithelial features, including desmosomes, hemidesmosomes, apical junctional complexes, and tissue-specific surface specializations. Proliferation and differentiation dynamics, crucial processes for epithelial homeostasis, are maintained in epithelioids reinforcing their self-sustaining capabilities. Functional assays indicated that the preserved architecture is physiologically active as human epithelioids formed a selective barrier and mounted robust wound-repair responses after injury. Together, these observations demonstrated that human epithelioids capture key functional features of native epithelia and therefore provide a more tissue-relevant setting to study clonal behavior.

The persistence of some *CD45*^+^/*CD3*^+^ cells in human esophageal epithelioids, together with comparable immune infiltration in several other human epithelioids after 1.5 - 2 months in continuous culture, suggests that certain levels of immune-epithelial original interactions can be maintained in these cultures. This is important because preservation of immune-epithelial crosstalk is a key requirement of novel culture systems^138–140^ and while a detailed identity and function of these immune populations remain to be clearly defined, their persistence may broaden the potential use of human epithelioids beyond the epithelial compartment alone.

An important question for any *in vitro* model is whether donor-derived clonal diversity is preserved or rapidly replaced by culture-driven selection^15,135,141,142^. Targeted NanoSeq showed that human epithelioids remain strongly polyclonal, with most individual cultures composed of large numbers of small clones and with occasional larger driver-bearing expansions. The spectrum of positively selected genes inferred from dN/dS also broadly recapitulated *in vivo* selection landscapes^6,40,118,119,143^, supporting the view that epithelioids retain key features of the donor’s clonal composition. This provides a tractable setting in which to examine how perturbations might reshape competition among pre-existing mutant lineages.

A methodological advantage of this platform compared with other culture methods is that epithelioids continuously shed differentiated cells, allowing liquid biopsy-like sampling of culture supernatants over time. Using targeted NanoSeq to assess the clonal composition of supernatants provided a non-invasive readout that was highly correlated with matched endpoint cultures. This overcomes the limitations of destructive sampling and would make it possible to resolve clone-specific dynamics and distinguish transient from persistent competitive responses during perturbations. These features may make epithelioids an attractive system to search for drugs capable of reducing the size or frequency of pre-tumourigenic clones, as a novel strategy for molecular cancer prevention.

By supporting prolonged epithelial maintenance and quantitative analysis across multiple human tissue types *in vitro*, human epithelioids expand the current repertoire of epithelial culture systems and provide a broadly applicable resource for studying clonal dynamics, somatic evolution, and tissue responses to genetic, environmental, and therapeutic perturbations in a human-relevant context^137^.

### Technical considerations

Some limitations of this work need to be considered. Epithelioids are epithelial-dominant cultures and therefore they might not fully capture stromal or immune interactions that can impose different levels of selection *in vivo*^147,148^. While we have demonstrated that human epithelioids recapitulate driver composition found *in vivo*, the establishment from small tissue fragments can bias the representation of different driver mutations in the resulting culture, particularly when clone sampling variance is high. In addition, supernatants might slightly over-represent differentiated or stressed cells, making an initial benchmarking against basal-enriched fractions important when the biological question specifically concerns stem or progenitor cell competition. Although these are relevant considerations and suggest opportunities for further optimization, they do not undermine the broad utility of human epithelioids for *in vitro* modelling of human epithelia, somatic selection, and preclinical testing of novel strategies for molecular cancer prevention.

## Methods

### 1. Human samples

Ethical approval for human tissue retrieval from deceased organ donors was obtained from the Cambridge South and Cambridge East Research Ethics Committees (Research Ethics Committee protocols 15/EE/0152 NRES Committee East of England-Cambridge South). Tissue was collected with the informed consent from the donors or when appropriate, their next-of-kin at Cambridge University Hospitals NHS Foundation Trust (Cambridge, UK). A segment of trachea (n=13), skin (n=9), buccal mucosa (n=13), esophagus (n=12), bladder (n=15), urethra (n=9) and endometrium (n=2), were resected within 60 minutes of circulatory arrest and preserved in ice-cold PBS buffer until arrival at the laboratory.

Ethical approval for submandibular gland tissue retrieval from head and neck squamous cell carcinoma patients undergoing neck dissection for radiologically confirmed nodal metastases was obtained from the North East- York Research Ethics Committee (Research Ethics Committee protocols 19/NE/0290, Substantial Amendment 12 - Collect sub-study). Tissue was obtained with patient’s informed consent at Cambridge University Hospitals NHS Foundation Trust, as part of the Hamlet.rt Collect study (Cambridge, UK). Submandibular gland biopsies (n=4) were preserved in complete FAD (cFAD) medium (see 2. Human epithelioid generation and maintenance) until arrival at the laboratory.

### 2. Human epithelioid generation and maintenance

Human epithelioids were generated and maintained using cFAD medium, containing Dulbecco’s modified Eagle’s medium (DMEM) (Gibco, 41966-029) and Nutrient Mixture F12 (DMEM/F12) (Gibco, 11320-074) at a ratio of 1:1. Medium was supplemented with 5% fetal bovine serum (FBS, Gibco, A5256701), 0.132 mM adenine (Sigma-Aldrich, A3159), 5 µg/ml insulin (Sigma-Aldrich, I5500), 5 µg/ml apo-transferrin (Sigma-Aldrich, T2036), 1% penicillin-streptomycin (Gibco, 15140-122), 1% amphotericin (Gibco, 15290-026), 0.1 nM cholera toxin (Sigma-Aldrich, C8052), 10 ng/ml epidermal growth factor (PeproTech EC, 100-15), 0.5 µg/ml hydrocortisone (Calbiochem, 386698).

Upon arrival at the laboratory, human samples were promptly washed in freshly prepared, ice-cold cFAD medium containing 12.5 µg/ml Plasmocin® (InvivoGen), 2% penicillin-streptomycin and 2% amphotericin for 30 minutes at 4°C. Plasmocin® was added to the cFAD medium during the first two weeks of culture to prevent potential donor-derived mycoplasma contamination. Samples were processed in sterile conditions under a dissecting scope. A small segment (∼5 mm × 5 mm) of each human sample was first reserved for optimal cutting temperature (OCT, VWR, 361603E) embedding and stored at -70°C for subsequent cryosectioning. Another small segment (∼ 3 mm × 3 mm) of each human sample was set aside for DNA isolation and mycoplasma testing by quantitative polymerase chain reaction (qPCR) using the PhoenixDx® Mycoplasma Mix (Procomcure Biotech, PCCSKU15209). Mycoplasma testing was carried out by the Research Instrumentation and Cell Services (RICS) at the CRUK Cambridge Institute, Cambridge, UK.

The remaining tissue was processed for epithelioid generation. The muscle layer and, where applicable, the majority of submucosa, were first removed following different strategies, depending on the epithelial tissue subtype:

- For trachea, bladder and urethra samples, the muscle layer was mechanically peeled off using fine forceps (SLS, INS1036).
- For skin, hair was first removed. Then, the attached fat, muscle, and excess of dermis were cut using fine scissors and scrapped off with a sterile scalpel (Swann-Morton™, 11798343).
- For buccal mucosa, the muscle layer and submucosa were cut and scraped off using fine scissors and a sterile scalpel.
- For esophagus, the muscle layer was peeled off using fine forceps and the epithelium and submucosa were initially cut using fine scissors. Any remaining excess of submucosa was removed using a sterile scalpel after processing tissue into smaller pieces.
- Submandibular gland was chopped into small pieces using a sterile scalpel without further dissection.
- For endometrium, the muscle layer was removed by scraping with a sterile scalpel.

Transwell ThinCert™ tissue culture inserts (Greiner Bio-One) were used for epithelioid generation and maintenance. Samples were cut into small explants (around 1 mm²) and plated onto 0.4µm pore-size ThinCert® cell culture inserts (Greiner Bio-One, 657641 [6-well] or 665641 [12-well]), depending on the experimental aim. Explants were plated with the epithelium facing upwards and the submucosa in contact with the insert membrane. A small drop of medium was maintained on top of each explant to keep the tissue moist and prevent desiccation. For 6-well inserts, 4 explants were placed on each insert, with 1 mL of cFAD medium added to the bottom compartment. For 12-well inserts, 2 explants and 0.5 mL of medium were used. Cultures were incubated at 37°C in 5% CO_2_ atmosphere.

Epithelial cell migration was typically observed ∼2-5 days after plating, forming cellular outgrowths. At that point, culture medium was topped up to the final working volumes for each compartment (6-well: 3 mL bottom and 1 mL top; 12-well: 1 mL bottom and 0.5 mL top) by adding fresh media to both compartments of the insert. Tissue explants were maintained within the epithelioids throughout their time in culture until downstream processing at the specified time points. Epithelial cells expanded until reaching the insert walls, achieving confluency within 2 to 3 weeks. Once the epithelioids reached confluency, medium from the top compartment was removed, and cultures were maintained at an air-liquid interface (ALI) for at least 2 weeks (or longer as indicated). Media were replenished twice weekly.

### 3. Immunofluorescence: Epithelioids and Cryosections

#### 3.1. Epithelioids

Epithelioids were fixed in 4% Paraformaldehyde (PFA) (FD NeuroTechnologies, INC. PF101) for 30 minutes at room temperature. Inserts were then washed 3 times in PHEM buffer (15 mM PIPES, 6.25 mM HEPES, 2.5 mM EGTA, 1 mM MgSO₄·7H₂O) for 10 minutes each. Skin epithelioids were peeled off from the membrane by incubation with a 50mM EDTA-PBS solution at the bottom compartment followed by a 30-minute incubation at 37°C prior to fixation steps.

After fixation, membranes were detached from the insert, and epithelioid biopsies were obtained by using 5 mm (KAI Medical, BP50F) or 7 mm punches (Acu-Punch, Acuderm Inc., P750). All subsequent steps were performed at room temperature, under gentle shaking. Membrane biopsies were transferred to 24-well plates (Greiner, 662102) and permeabilized in 1% PB buffer (0.5% bovine serum albumin [BSA; Sigma-Aldrich, A9647-100G], 0.25% fish skin gelatine [Sigma-Aldrich, G7765-1L], 1% Triton X-100 [Sigma-Aldrich, X100-500ML] in PHEM buffer) for 30 minutes. Membrane biopsies were then blocked in 1% Blocking buffer (0.5% BSA, 0.25% fish skin gelatin, 1% Triton X-100 and 10% Donkey serum [Bio-Rad, C06SB] in PHEM buffer) for one hour. Incubation with primary antibodies diluted in 1% Blocking buffer was performed overnight. Membranes were then washed five times with PHEM buffer for at least 10 minutes. The following incubation with secondary antibodies diluted in 1% Blocking buffer was performed overnight. Cell nuclei were counterstained with 4′,6-diamidino-2-phenylindole (DAPI) (3 µg/mL, Sigma-Aldrich, D9542). Where specified, Alexa Fluor^TM^ 488 Phalloidin (1:1000, Invitrogen A12379) or Alexa Fluor^TM^ 647 anti-human/mouse CD49f (1:500; BioLegend, 313610) were also included in the secondary antibody solution. Primary and secondary antibodies, references and dilutions are summarized in **Supplementary table 1**. Samples were washed five times in PHEM Buffer for 10 minutes each and mounted on glass microscopy slides (Epredia, AG00000112E01MNZ10) using either a PBS-Glycerol (Sigma-Aldrich, G7757- 1L) solution (ratio 1:1) or RapiClear® (SunJin Lab, RC152002) to enable deep imaging for thicker epithelioids (>50 µm). Samples were covered with glass coverslips (VWR, 631-0136), sealed with nail polish and air-dried overnight.

#### 3.2. Cryosections

OCT-embedded tissues were mounted on a Cryostat (Leica, CM3050 S) and sectioned at 10 μm onto glass microscopy slides (Superfrost^TM^ Plus Adhesion Microscopy slides, Epredia, J1800AMNZ). Slides were air dried for 30 minutes at room temperature and fixed in 4% PFA for 5 minutes at room temperature. Next, tissues were rehydrated by immersion in PHEM buffer twice for five minutes each. Blocking stage was performed covering the slices with 0.5% Blocking buffer (0.5% BSA, 0.25% fish skin gelatin, 0.5% Triton X-100 and 10% Donkey serum in PHEM Buffer) for 30 minutes at room temperature inside of a humidified StainTray^TM^ (Simport, M918). Slides were then stained using primary antibodies diluted in 0.5% Blocking Buffer and incubated overnight at room temperature inside a StainTray^TM^ chamber, followed by five washes with PHEM buffer, for 5 minutes each, at room temperature. The following incubation with secondary antibodies diluted in 0.5% Blocking buffer was performed for 3 hours at room temperature inside a StainTray^TM^ chamber. For nuclear visualization, 3 µg/mL of DAPI (Sigma-Aldrich, D9542) were added to the solution containing secondary antibodies. When specified, Alexa Fluor^TM^ 488 phalloidin (1:1000) or Alexa Fluor^TM^ 647 anti-human/mouse CD49f (1:500) were also included in the secondary antibody solution. Primary and secondary antibodies, references and dilutions are summarized in **Supplementary table 1.** After incubation with secondary antibodies, slides were washed with PHEM buffer five times, for 5 minutes each, at room temperature. Finally, samples were mounted using a Glycerol-PBS 1:1 solution.

#### 3.3. Image and image analysis

Confocal imaging was performed using a Nikon Ti2-E inverted microscope equipped with a CSU-W1 SoRa spinning disk confocal unit, operated with NIS-Elements Acquisition software (version AR 5.42.06).

Epithelioids were imaged using a 40x Plan water immersion objective, with a numeric aperture (NA) of 1.15 and a long working distance (LWD) of 0.61 mm (Nikon). 3D z-stacks were generated by sequential imaging at 0.3 µm z-steps across the full thickness of the epithelioids.

Rendered 3D z-stacks and orthogonal views were created using Volocity v.6.3 (Perkin Elmer). Single plane views were generated using either Volocity v.6.3 (Perkin Elmer) or Fiji^1^.

### 4. EdU labelling for proliferation and differentiation

Proliferation and differentiation were measured in human epithelioids five weeks after the generation of the culture. cFAD medium containing 10µM of EdU (5-ethynyl-2′-deoxyuridine, Invitrogen, C10338) was added to the bottom compartment of the transwell plate. Cultures were incubated for 2-8 hours at 37°C in a humidified 5% CO₂ incubator. Following the pulse, EdU-containing medium was removed and replaced with fresh cFAD. Cultures were then incubated at 37°C in 5% CO₂ for an additional 1 hour for proliferation analysis, or 48-96 hours for differentiation analysis. At the indicated time points, cultures were fixed in 4% paraformaldehyde (PFA) for 30 minutes at room temperature. EdU detection was performed using the Click-iT EdU imaging kit (Invitrogen, C10338) according to the manufacturer’s instructions.

For proliferation and differentiation analysis using EdU labelling, epithelioids were scanned using a 20x Plan-Apochromat water immersion objective, with a NA of 0.95 and WD of 0.95 mm (Nikon). 3D z-stacks for differentiation analysis were generated by sequential imaging at 0.6 µm z-steps across the full thickness of the epithelioids. A minimum of 9 fields of view were collected for this analysis. Representative images were acquired using a 40x Plan water immersion objective, with a NA of 1.15 and a long WD (LWD) of 0.61 mm (Nikon).

Basal cell density (nuclei) and cell proliferation and differentiation (EdU) were quantified using the DSBDancer v8.0.2, a plugin for Fiji developed at the Gurdon Institute Imaging Facility (GIIF). For EdU analysis, the EdU channel was used to select the best-focused slice from each z-stack. Nuclei were segmented in the frequency-filtered DAPI channel using Otsu’s method (Otsu 1979, IEEE Transactions on Systems, Man and Cybenetics), with a radius range of 2.5-5 µm. Nuclei were classified as EdU-positive if their mean EdU intensity had a Z-score of 0.6 standard deviation or higher. Cell differentiation (suprabasal EdU) was assessed using a semi-automated approach that combined the DSBDancer pipeline with manual quantification of EdU-positive cells in the suprabasal layers of the epithelioids. Basal cell density was calculated as the number of segmented nuclei per mm^2^.

**Supplementary table 1.** Primary and secondary antibodies.

| Antibody | Target/protein | Identifier | Dilution | Supplier |
| --- | --- | --- | --- | --- |
| Primary | CD3ε | Cat# ab5690,<br>RRID:AB_305055 | 1 in 200 | Abcam Ltd |
| Primary | CD45 | Cat# 368502,<br>RRID:AB_2566240 | 1 in 400 | BioLegend |
| Primary | CD49f | Cat# 313602<br>RRID:AB_345296 | 1 in 200 | BioLegend |
| Primary | CD49f<br>Alexa Fluor™ 647 | Cat# 313610<br>RRID:AB_493636 | 1 in 500 | BioLegend |
| Primary | E-Cadherin | Cat# #3195<br>RRID:AB_2291471 | 1 in 1000 | Cell Signaling |
| Primary | FOXA2 | Cat# ab108396<br>RRID:AB_10863255 | 1 in 200 | Abcam Ltd |
| Primary | FOXJ1 | Cat# 14-9965-82,<br>RRID:AB_1548835 | 1 in 1000 | Thermo Fisher Scientific |
| Primary | Involucrin | Cat# MA5-11803,<br>RRID:AB_10982738 | 1 in 1000 | Thermo Fisher Scientific |
| Primary | KRT4 | Cat# C5176,<br>RRID:AB_258888 | 1 in 500 | Sigma-Aldrich |
| Primary | KRT5 | Cat# 905901,<br>RRID:AB_2565054 | 1 in 500 | BioLegend |
| Primary | KRT7 | Cat# MA5-11986,<br>RRID:AB_10989596 | 1 in 500 | Thermo Fisher Scientific |
| Primary | KRT8 | Cat# ab53280,<br>RRID:AB_869901 | 1 in 1000 | Abcam Ltd |
| Primary | KRT10 | Cat# ab9025,<br>RRID:AB_2134556 | 1 in 500 | Abcam Ltd |
| Primary | KRT13 | Cat# BP5076,<br>RRID:AB_979608 | 1 in 500 | OriGene EU |
| Primary | KRT14 | Cat# 905301,<br>RRID:AB_2565048 | 1 in 1500 | BioLegend |
| Primary | KRT20 | Cat# GP-K20<br>AB_1541036 | 1 in 200 | Progen |
| Primary | Ki67 | Cat# ab15580,<br>RRID:AB_443209 | 1 in 1000 | Abcam Ltd |
| Primary | KLF4 | Cat# AF3158,<br>RRID:AB_2130245 | 1 in 100 | R&D Systems |
| Primary | MIST1 | Cat# ab187978,<br>RRID:AB_2924393 | 1 in 100 | Abcam Ltd |
| Primary | MUC5AC | Cat# MA1-38223,<br>RRID:AB_2266697 | 1 in 500 | Thermo Fisher Scientific |
| Primary | Anti-MUC5B | Cat# ab77995,<br>RRID:AB_2146987 | 1 in 200 | Abcam Ltd |
| Primary | SCGB1A1 | Cat# 10490-1-AP,<br>RRID:AB_2183285 | 1 in 500 | Proteintech |
| Primary | SOX2 | Cat# AF2018,<br>RRID:AB_355110 | 1 in 250 | R&D Systems |
| Primary | TP63 | Cat# GTX102425, | 1 in 1000 | GeneTex |
|  |  | RRID:AB_1952344 |  |  |
| Primary | Acetylated tubulin | Cat# T6793, RRID:AB_477585 | 1 in 1000 | Sigma-Aldrich |
| Secondary | Goat anti-Chicken IgY, Alexa Fluor™ 488 | Cat# A-11039, RRID:AB_2534096 | 1 in 1000 | Fisher Scientific |
| Secondary | Donkey anti-Goat, Alexa Fluor™ Plus 555 | Cat# A32816, RRID:AB_2762839 | 1 in 3000 | Fisher Scientific |
| Secondary | Goat anti-Guinea Pig, Alexa Fluor™ 647 | Cat# A-21450, RRID:AB_2535867 | 1 in 1000 | Fisher Scientific |
| Secondary | Donkey anti-Mouse, Alexa Fluor™ Plus 488 | Cat# A32766, RRID:AB_2866493 | 1 in 3000 | Fisher Scientific |
| Secondary | Donkey anti-Mouse, Alexa Fluor™ Plus 555 | Cat# A32773, RRID:AB_2762848 | 1 in 3000 | Fisher Scientific |
| Secondary | Donkey anti-Mouse, Alexa Fluor™ Plus 647 | Cat# A32787, RRID:AB_2866494 | 1 in 3000 | Fisher Scientific |
| Secondary | Donkey anti-Rabbit, Alexa Fluor™ Plus 488 | Cat# A32790, RRID:AB_2866495 | 1 in 3000 | Fisher Scientific |
| Secondary | Donkey anti-Rabbit, Alexa Fluor™ Plus 555 | Cat# A32794, RRID:AB_2762834 | 1 in 3000 | Fisher Scientific |
| Secondary | Donkey anti-Rabbit, Alexa Fluor™ Plus 647 | Cat# A32795, RRID:AB_2866496 | 1 in 3000 | Fisher Scientific |
| Secondary | Donkey anti-Rat, Alexa Fluor™ Plus 555 | Cat# A48270, RRID:AB_2896336 | 1 in 3000 | Fisher Scientific |

### 5. Electron microscopy

Membrane biopsies from human epithelioid cultures were obtained by using a 5 mm diameter biopsy punch, and fixed in a fixative solution containing 4% PFA (Sigma-Aldrich, P6148-500G) and 2.5% glutaraldehyde (EM grade distillation purified, Electron Microscopy Sciences, 16210) in 0.1 M phosphate buffer pH 7.4 (PB) (0.2 M Na_2_HPO_4_, Acros Organics, 448150010; 54 mM NaH_2_PO_4_, Acros Organics, 447760010; dissolved in ddH_2_O, pH 7.4) at 37°C for 15 minutes. The fixative was refreshed, and samples were incubated for a further 2 hours at 4°C, followed by a final incubation with fresh fixative at 4°C for 72 hours.

#### 5.1 Transmission electron microscopy (TEM)

Sample processing was performed at the Electron Microscopy Facility, Wellcome-MRC Cambridge Stem Cell Institute, Cambridge, UK. First, samples were washed three times with 0.22 µm-filtered dH_2_O for 10 minutes. Post-fixation was carried out in 1% osmium tetroxide (TAAB Laboratories Equipment Ltd, TAAB OO12) with 1.5% potassium ferrocyanide (Scientific Laboratory Supplies Ltd, CHE2956) for 30 minutes, followed by additional dH2O washes. Samples were then stained overnight at 4°C using an 80% solution of UA-Zero (Agar Scientific, AGR1000) diluted in ethanol and washed again in dH₂O. Dehydration was performed in an ethanol series (70%, 95%, 100%), with 10 minutes at each concentration under gentle shaking, followed by an additional incubation in 100% ethanol for 10 minutes. For epoxy resin embedding, samples were transferred to flat embedding molds using forceps, oriented with the cell-containing side of the transwell insert facing up. Molds were filled with fresh resin (Agar Scientific, Agar 100 Resin Kit) and polymerized at 60 °C for 60 hours.

Ultrathin sections (∼80 nm) were cut with an Ultracut UC-6 (Leica Biosystems), collected on Formvar-coated single-slot copper grids (Sigma, 930288), stained with lead citrate (Reynold’s solution). All images were taken with a Xarosa digital camera (EMSIS GmbH, Münster, Germany) controlled by Radius software (Version 2.1) at the Electron Microscopy Core Facility, Centro de Investigación Príncipe Felipe (CIPF) (Valencia, Spain).

#### 5.2 Scanning electron microscopy (SEM)

Sample processing was performed at the Electron Microscopy Facility, Wellcome-MRC Cambridge Stem Cell Institute, Cambridge, UK. Samples were washed twice with 0.22 µm-filtered dH₂O for 5 minutes each. Dehydration was carried out through an ethanol series (25%, 50%, 70%, 90%, and 100%) with incubation times of 5, 10, 30, 30, and 10 minutes, respectively, followed by two additional 10-minutes washes in 100% ethanol. Fresh 100% ethanol was added prior to loading the samples into the Critical Point Dryer (CPD). Within the CPD, 15 slow CO₂ exchange cycles were performed to gradually and completely replace ethanol with liquid CO₂. Samples were then mounted on aluminum stubs using double-sided carbon adhesive tabs, oriented with the cell-containing side of the insert facing up. Cell surface was analyzed using a SEM Inspect F50 (Thermo Fisher Scientific) at an acceleration voltage of 5 - 10 kV, at the Advanced Microscopy Laboratory, (Universidad de Zaragoza, Zaragoza, Spain).

### 6. Single-cell RNA sequencing and analysis

Human epithelioids were cultured for at least 3 weeks with media in the top compartment and then changed to ALI condition for 2 additional weeks. Cultures were washed twice with PBS and dissociated by subsequent incubation with pre-warmed TrypLE™ Express Enzyme (Gibco, 12604021) (2 mL at the bottom and 0.5 mL at the top of each insert) at 37°C for 10 minutes, followed by gentle insert scrapping with a pipette tip to facilitate cell detachment. This step was repeated according to the specific requirements of each epithelioid type for up to 50 minutes. To maximize cell viability, all subsequent steps were carried out on ice, and all solutions were ice-cold. Dissociated cells were transferred to individual tubes containing 7 mL of cFAD to inactivate TrypLE™ and centrifuged at 400 g for 5 minutes at 4°C. Cells were washed with cFAD and centrifuged again. Cell pellets were resuspended in 2 mL of PBS + 0.04% BSA to reduce cell aggregation and filtered through a 50 µm pore-size cell strainer (CellTrics®, 04-004-2327) to remove any clumps. An additional 3 mL of PBS + 0.04% BSA was added, and cell suspensions were homogenized by gentle up-and-down pipetting. Subsequently, 0.1 mL of each suspension was mixed 1:1 with a 0.4% Trypan Blue solution (Gibco, 15250061) for cell counting and viability assessment using a counting chamber (Hirschmann, 8100103). A final centrifugation step was carried out, and cells were resuspended in 0.2–1 mL of PBS + 0.04% BSA, yielding a final concentration of 1–2 × 10⁶ cells/mL

Cells were loaded into the 10x Genomics Chromium controller according to the manufacturer’s protocol using a Chromium Next GEM Single Cell 3′ v3.1 four-reaction kit (10x Genomics, PN-1000269). Libraries were prepared following the manufacturer’s instructions, with a target recovery of 7,000 cells per reaction. Sequencing was performed on an Illumina NovaSeq 6000 platform at the Wellcome Sanger Institute Single Cell Genomics Core facility, Hinxton, UK, to an average depth of 604,578,365 reads per library.

Raw base call files were processed with Cell Ranger software (v8.0, https://support.10xgenomics.com/single-cell-gene-expression) using the GRCh38-2024-1 human reference genome. Read count matrices were analyzed in Python using Scanpy (1.11.4) following standard workflows. Cell cycles scores were calculated using scanpy.tl.score_genes_cell_cycle function with a published gene list^2^. For gene score calculation, the analysis followed a previously described procedure^3^. The top 100 most highly expressed genes from each cluster were used to define a gene set, and scanpy.tl.gene_score function was used to score the expression of these sets in our data.

**Supplementary table 2.** Software and algorithms used for single-cell RNA sequencing analysis.

| Software and algorithms |  |  |
| --- | --- | --- |
| Fiji | Schindelin et al.,<br>2012 | https://fiji.sc/ |
| **Python (v3.11.10)** | - | https://www.python.org/ |
| **Cell Ranger Single-Cell Software Suite (v8.0)** | - | https://www.10xgenomics.com/support/software/cell-ranger/downloads |
| **Scanpy (1.11.4)** | Wolf et al., 2018 | https://scanpy.readthedocs.io/en/stable/ |

### 7. Barrier formation assay

Human epithelioids derived from trachea, skin, buccal mucosa, esophagus, and bladder were grown in 12-well inserts and changed to ALI 1 week prior to the permeability assays. Epithelioids were washed once with PBS and incubated for 1 hour with 4 KDa fluorescein isothiocyanate-dextran (Dextran-FITC, 50 µg/mL) (Sigma-Aldrich, 46944) in cFAD prepared with FluoroBrite™ DMEM (Gibco, A1896701) and DMEM/F12, no phenol red (Gibco, 21041025) (1:1). 0.5 mL of Dextran-FITC solution was added to the top of each insert while 1 mL of Fluorobrite cFAD medium was added to the bottom. At the end of the incubation time, the bottom medium was homogenized by gentle shaking, and 0.1 mL was collected and transferred onto a flat-bottom white 96 well-plate. Fluorescence intensity was measured in a plate reader (CLARIOstar, BMG LABTECH) using the following excitation/emission filters: 482-10/516-8. Fresh Fluorobrite™ medium was used as blank control. An empty insert with 4 KDa Dextran-FITC added to the top was used as positive control for maximum permeability. The fluorescence intensity of each epithelioid sample was normalized to the corresponding positive control.

### 8. Liquid biopsy collection

The collection of liquid biopsies from ALI-cultured human epithelioids was performed twice a week. For collection, 1 mL of PBS was added to the top of the insert, and plates were then incubated for 10 minutes at 37°C in a 5% CO_2_ atmosphere. Following incubation, inserts were washed by pipetting PBS up and down to boost cell shedding and maximize sample collection. PBS containing shed cells was transferred to a sterile 1.5 mL tube. Cells were then pelleted by centrifugation (600 g for 5 minutes). The same tube was used for subsequent collections. The number of collections ranged from 8 to 10. Tubes containing cell pellets were kept at 4°C until the last collection. Once the last collection was completed, cells were pelleted and resuspended in 180 µL of ATL tissue lysis buffer (from the QIAamp DNA Micro Kit, Qiagen, 56304).

Epithelioid cultures were digested after the last supernatant collection for comparison of mutational landscapes in the cultures and the supernatants. Prior to lysis, remaining explants were removed and 180µL or 360µL ATL tissue lysis buffer was added to the insert, and cells were scraped and mixed until the obtention of a homogeneous lysate.

### 9. DNA isolation and quantification

DNA isolation from epithelioids and tissue biopsies was performed using the DNeasy Blood and Tissue Kit (Qiagen, 69506). DNA isolation from supernatants was carried out using the QIAamp DNA Micro Kit (Qiagen, 56304). The protocol was followed according to the manufacturer’s instructions, with minor modifications to maximise DNA yield and remove contaminants. Specifically, before elution, columns were left open for 15-20 minutes to allow residual ethanol to evaporate. The elution buffer (Buffer AE) was pre-warmed to 70°C, applied to the column, and incubated for 3 minutes at room temperature before elution. Last, the eluate was re-transferred to the column and eluted again.

DNA concentration was measured using a Qubit™ 4 Fluorometer (ThermoFisher, Q33226) and the Qubit™ 1X dsDNA Broad Range Kit (ThermoFisher, Q33265), following the manufacturer’s instructions.

### 10. Wound healing and regeneration in epithelioids

2 months-old epithelioids were wounded under sterile conditions in two sequential steps. First, an 8 mm sterile biopsy punch (KAI Medical, BP-80F) was used to demarcate a circular wound area at the center of each epithelioid; after which cells within the wound area were removed using a micro scalpel blade (Carl Martin GmbH, 871MB/64). Epithelioids were then washed twice with cFAD medium to remove cellular debris.Brightfield whole-well tile scan images were acquired at specific time points using a Nikon Ti2-E inverted microscope equipped with a CSU-W1 SoRa spinning disk confocal unit and operated with NIS-Elements software (version AR 5.42.06). Images were acquired using a Prime 95B 1 camera and LED transmitted-light illumination. Wounded epithelioids were live-imaged using a 10x Plan-Apochromat objective (NA of 0.45 and a WD of 4.0 mm; Nikon). For EdU quantification, whole epithelioids were scanned using the same 10x Plan-Apochromat objective. Representative images of regions both adjacent to and distant from the wound area were acquired using a 20x Plan-Apochromat water immersion objective (NA 0.95, WD 0.95 mm; Nikon). Epithelioid wound healing was quantified in Fiji^1^ by manually defining a region of interest (ROI) corresponding to the wound area in each time-lapse image. ROIs excluded areas covered by cells, capturing only the cell-free wound space. EdU positive cell counts were then quantified using the DSBDancer plugin in Fiji (see Section 3.2.3).

### 11. Duplex sequencing: Targeted Nanoseq

A total of 237 NanoSeq libraries were prepared as described in Lawson et al, using a targeted panel of 239 genes (Appendix A1, Table A.1), and sequenced on NovaSeq X instruments. These libraries corresponded to 17 original tissue samples from 9 donors, including skin (6 epithelioids/1 donor), buccal mucosa (24/4), esophagus (26/5), trachea (24/4), and bladder (24/4), i.e. 104 epithelioids in total. For each epithelioid, supernatant cells and wells were sequenced separately. Libraries were sequenced to a median duplex coverage of 257 dx per library (range = 31-698 dx).

Variant calling was done with the NanoSeq pipeline (https://github.com/cancerit/NanoSeq) as in Lawson et al 2025, with the exception that here, when available, the original tissue was used as the matched normal.

Selection analyses were done with R package dndscv (https://github.com/im3sanger/dndscv), accounting for variation in the mean duplex coverage of each gene. Mutated cell fractions were estimated on genes under positive selection as the sum of duplex VAFs, with the lower bound estimate assuming mutations happen in cells as double hits, and the upper bound assuming they happen as single hits (2x duplex VAF).

Unbiased VAFs (see Lawson et al 2025) were calculated with the NanoSeq pipeline and were used to compare clonality between supernatant (mostly differentiated) and well cells (mix of basal and differentiated cells). When one of the two, supernatant or well, did not have the mutation called, then we used bam2R (from R package deepSNV) to genotype the mutation in the deduplicated bam file, using the following parameters: q = 30, mask = 3844, mq = 30.

Signature extraction for substitutions was done with R package sigfit (https://github.com/kgori/sigfit). We explored extractions between 3 and 6 signatures. Based on goodness of fit and visual inspection, we chose the 4 signatures solution as the optimal one. Extraction was conducted using 5000 iterations, 1000 of which for warm up. Exposure of each signature in each sample was done with 2000 iterations, 1000 for warm up. Exposures were only estimated on wells. Cosine similarities between these four signatures and known signatures were estimated using COSMIC 3.2

| donor | tissue | #tissue | #supernat. | #wells | #replicates |
| --- | --- | --- | --- | --- | --- |
| 900 | Bladder | 1 | 4 | 4 | 4 |
| 900 | Bladder | 0 | 24 | 7 | 7 |
| 898 | Bladder | 1 | 4 | 4 | 4 |
| 901 | Bladder | 1 | 3 | 3 | 3 |
| 906 | Bladder | 1 | 6 | 6 | 6 |
| 889 | Buccal mucosa | 1 | 6 | 6 | 6 |
| 890 | Buccal mucosa | 1 | 6 | 6 | 6 |
| 901 | Buccal mucosa | 1 | 6 | 6 | 6 |
| 907 | Buccal mucosa | 1 | 6 | 6 | 6 |
| 888 | Esophagus | 1 | 6 | 6 | 6 |
| 900 | Esophagus | 1 | 5 | 6 | 6 |
| 901 | Esophagus | 1 | 6 | 6 | 6 |
| 906 | Esophagus | 1 | 6 | 6 | 6 |
| 916 | Esophagus | 0 | 0 | 2 | 2 |
| 888 | Skin | 1 | 5 | 6 | 6 |
| 900 | Trachea | 1 | 5 | 6 | 6 |
| 898 | Trachea | 1 | 6 | 6 | 6 |
| 901 | Trachea | 1 | 6 | 6 | 6 |
| 906 | Trachea | 1 | 6 | 6 | 6 |

### 12. Statistical analysis

A minimum of n = 3 biological replicates was used for all statistical analyses, with exact replicate numbers reported in each figure legend. Quantitative results are presented as mean ± SD or mean, as specified in the corresponding figures. Graphs and analyses were generated using GraphPad Prism 10.6.1, Scanpy 1.11.4^4^ or R. For datasets with more than two groups, one-way ANOVA followed by Tukey’s or Dunnett’s multiple-comparisons post-hoc tests was applied. Confocal images are representative of at least n = 3 biological replicates, unless otherwise indicated in the figure legends.

---

**Figure S1:**
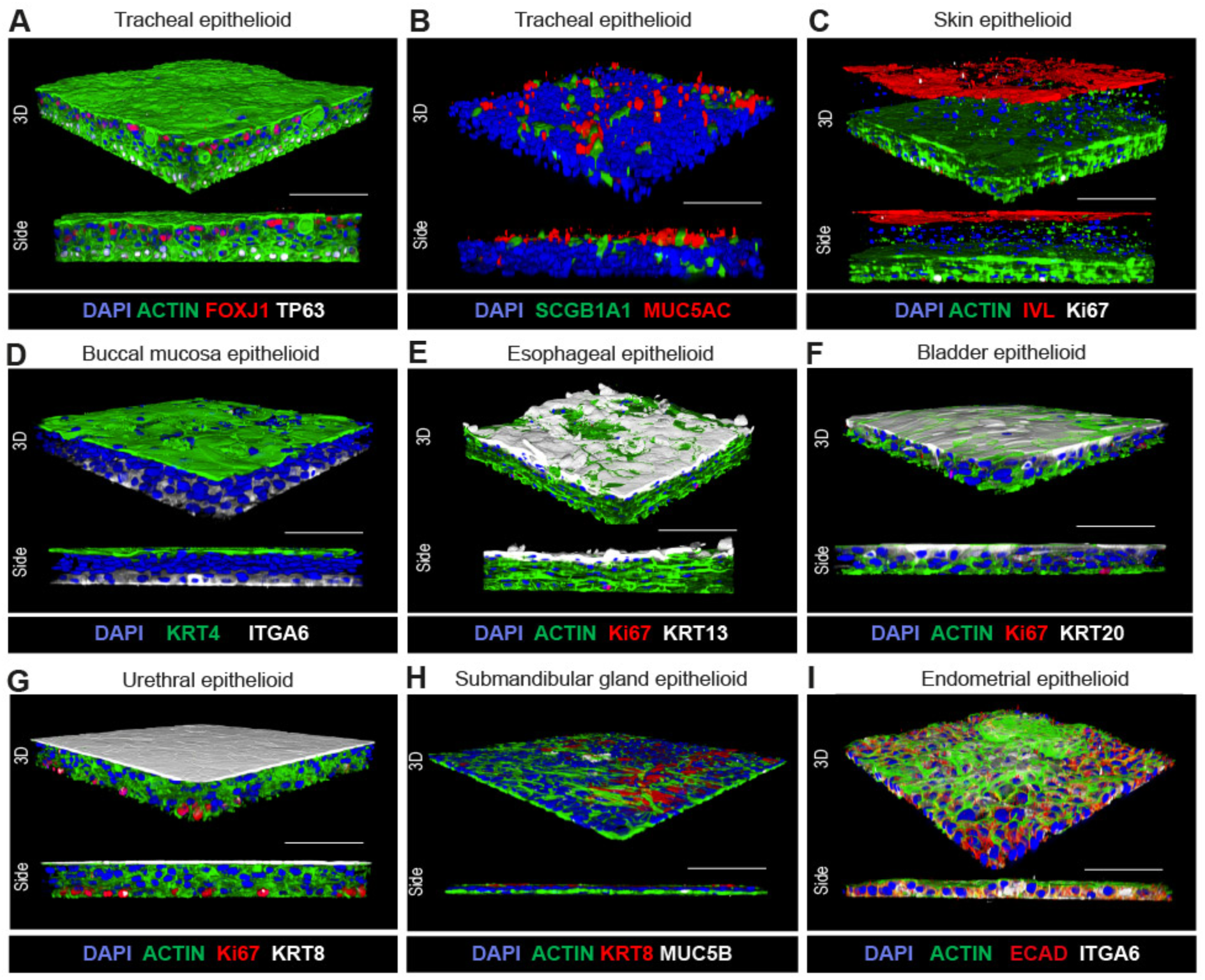
Extended data, Structural and phenotypic characterization of human epithelioids generated from a diversity of epithelia. (A –. **I)** Rendered 3D confocal images including different staining combinations to show the architecture and cell populations of human trachea **(A – B)**, skin **(C)**, buccal mucosa **(D)**, esophageal **(E)**, bladder **(F)**, urethral **(G)**, submandibular gland **(H)** and endometrial **(I)** epithelioids. Scale bar, 100 μm.

**Figure S2:**
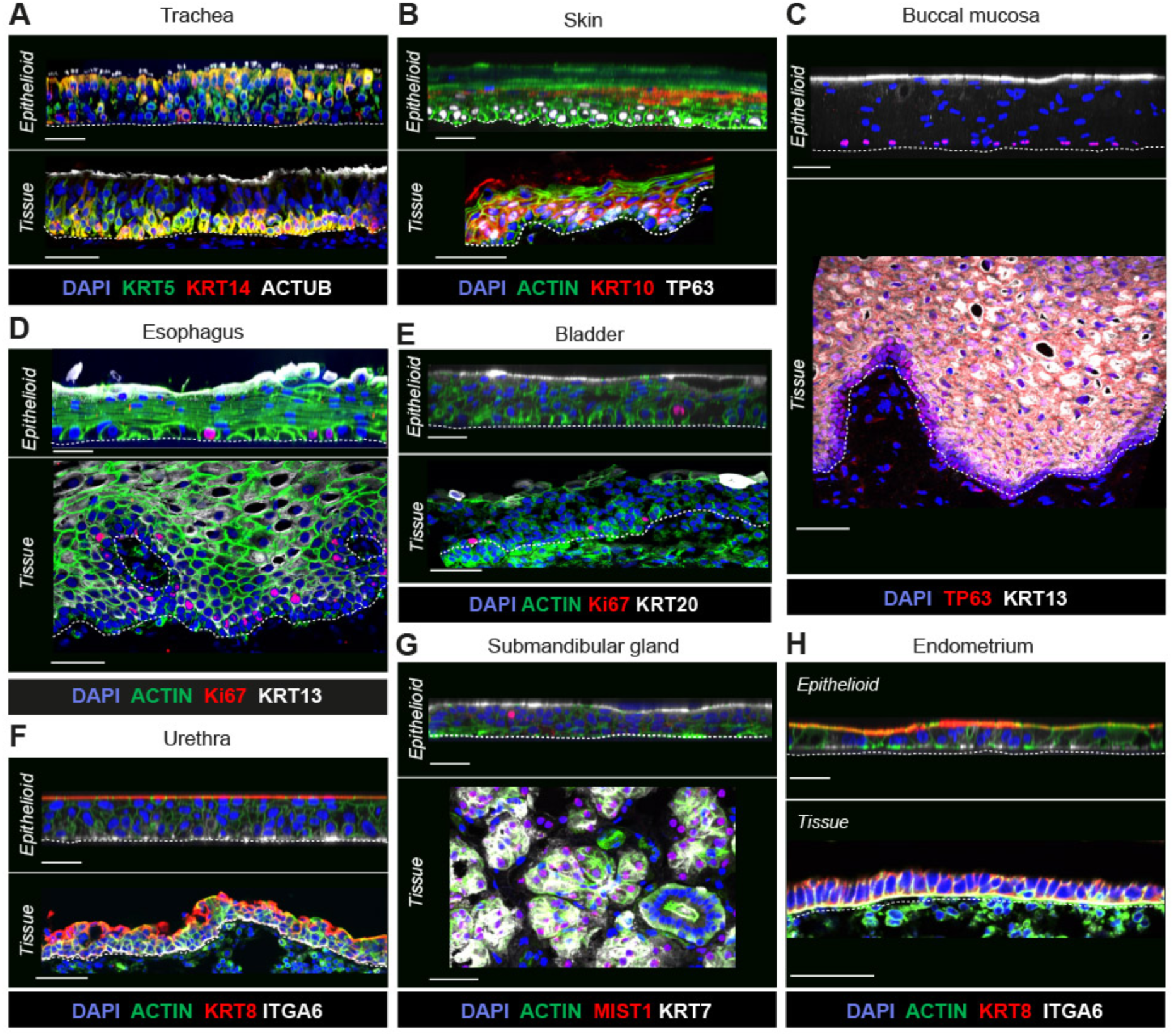
Extended data, Human epithelioids recapitulate the structure and cellular distribution of human epithelia. (A –. **H)** Images showing side views from 3D reconstructions of human tracheal **(A)**, skin **(B)**, buccal mucosa **(C)**, esophageal **(D)**, bladder **(E)**, urethral **(F)**, submandibular gland **(G)** and endometrial **(H)** epithelioids benchmarked against tissue cryosections. The dotted line denotes epithelioids or epithelia basal boundaries. Scale bar, 50 μm.

**Figure S3:**
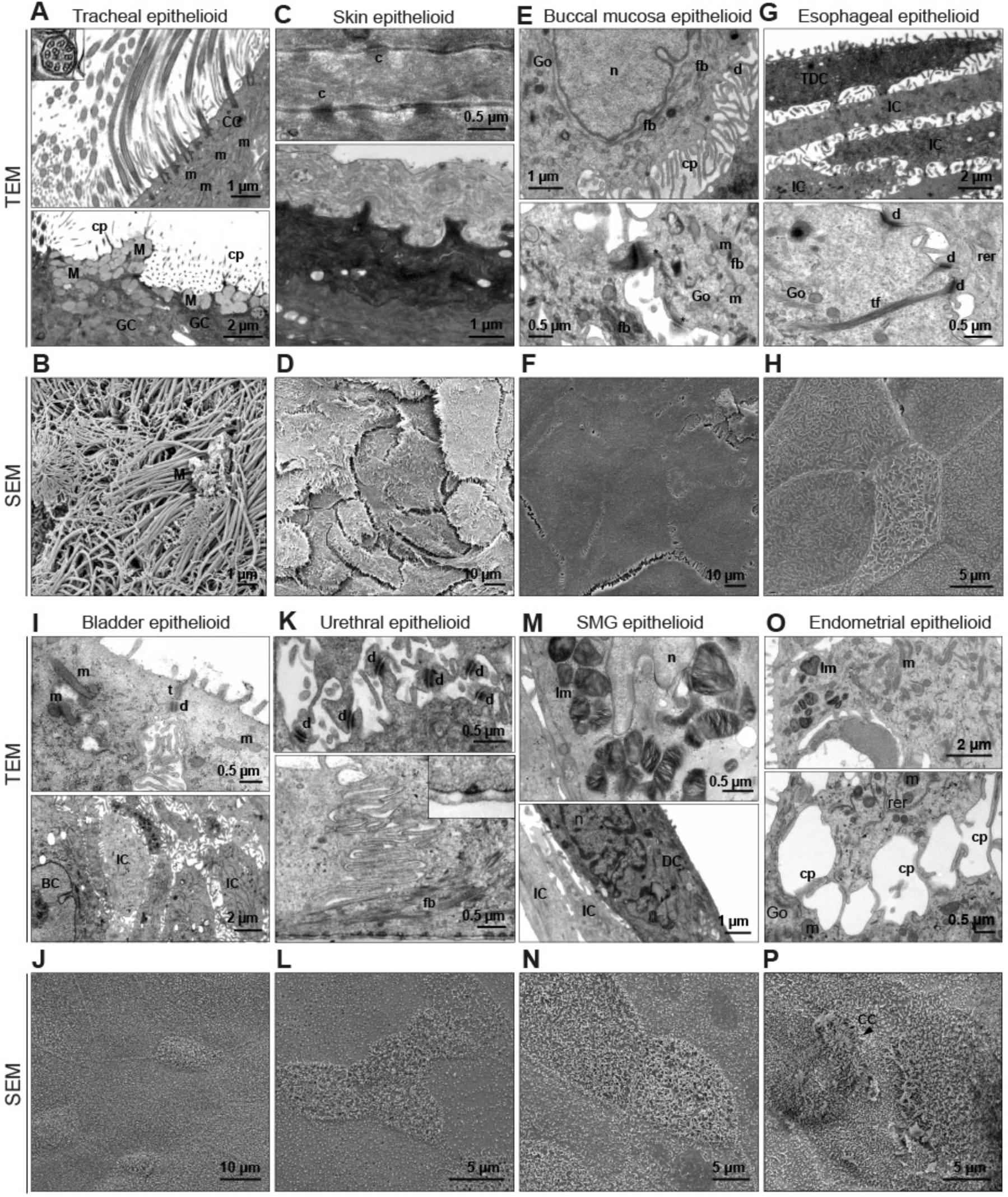
Extended data, Ultrastructural analysis of human epithelioids reveals apical-to-basal organization and epithelial integrity. (A –. **P)** Detailed analyses of epithelioids by TEM (top) and SEM (bottom) reveal the presence of specific subcellular structures and organelles in tracheal **(A – B)**, skin **(C – D)**, buccal mucosa **(E – F)**, esophageal **(G – H)**, bladder **(I – J)**, urethral **(K – L)**, submandibular gland **(M – N)** and endometrial **(O – P)** epithelioids. (BC: basal cells; CC: ciliated cells; cp: cytoplasmic prolongations; d: desmosome; fb: filament bundles; IC: intermediate cells; GC: globet cells; Go: Golgi apparatus; c: corneosomes; lb: lamellar bodies; m: mitochondria; M: mucus n: nucleus; rer: rough endoplasmic reticulum; TDC: terminally differenciated cells; tf: tonofilaments; t: tight junction).

**Figure S4:**
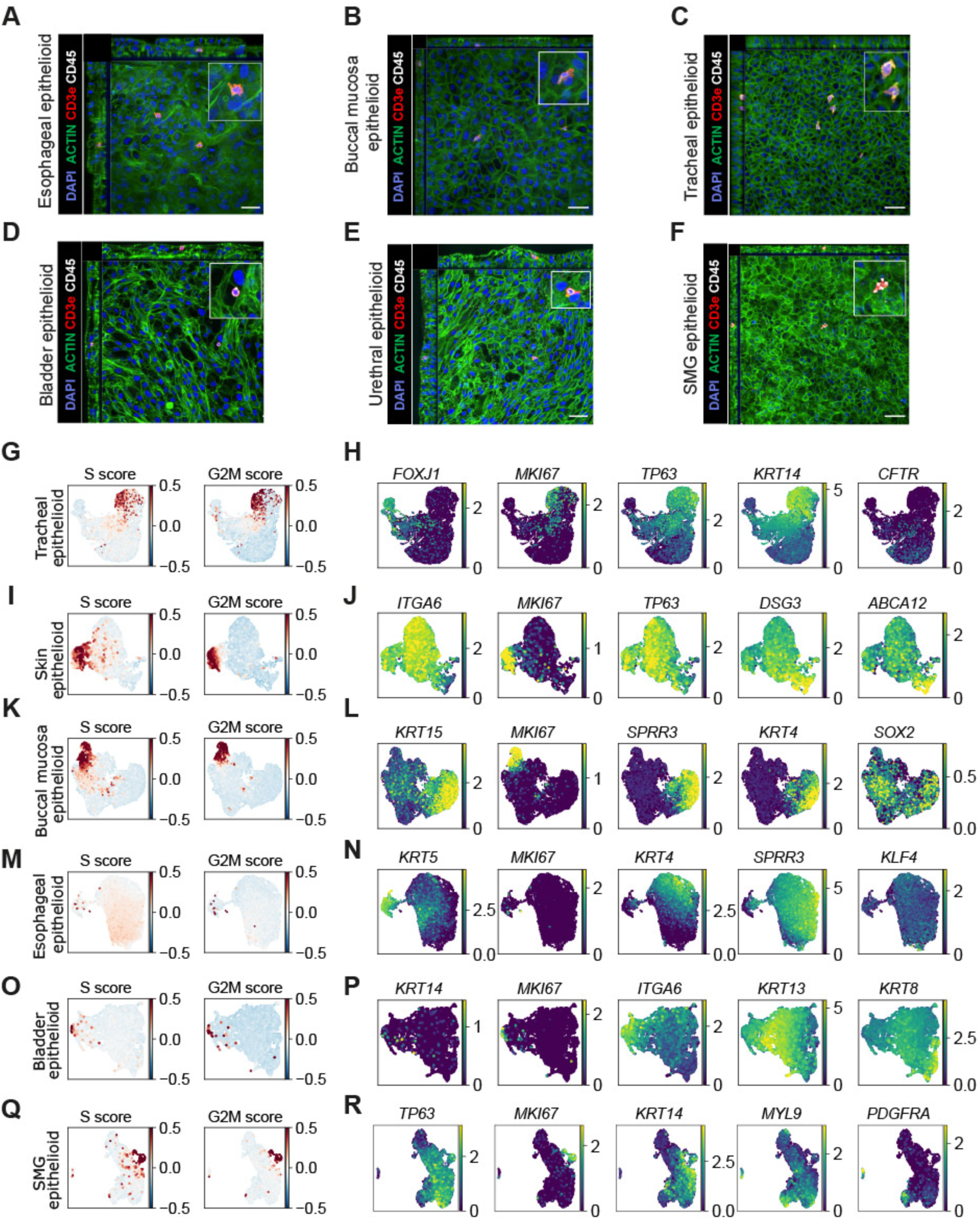
Extended data; Single-cell transcriptomics confirm the protein-level data and expose tissue-resident immune cells in human epithelioids. **(A – F)** Confocal images showing orthogonal reconstruction of human esophageal (A), buccal mucosa (B), tracheal (C), bladder (D), urethral (E) and submandibular gland (F) epithelioids displaying tissue-resident T cells, dually stained for CD45 (red) and CD3ε (white). Scale bar, 50 μm. **(G, I, K, M, O, Q)** UMAP plots showing cell cycle score cell score expression patterns across cell clusters, calculated from a published gene list (Kowalczyk et al., 2015). **(H, J, L, N, P, WR)** UMAP plots showing normalized expression (counts per 10,000 reads; CP10K) of selected marker genes in tracheal **(B)**, skin **(E)**, buccal mucosa **(H)**, esophageal **(K)**, bladder **(N)** and submandibular gland **(Q)** epithelioids.

**Figure S5:**
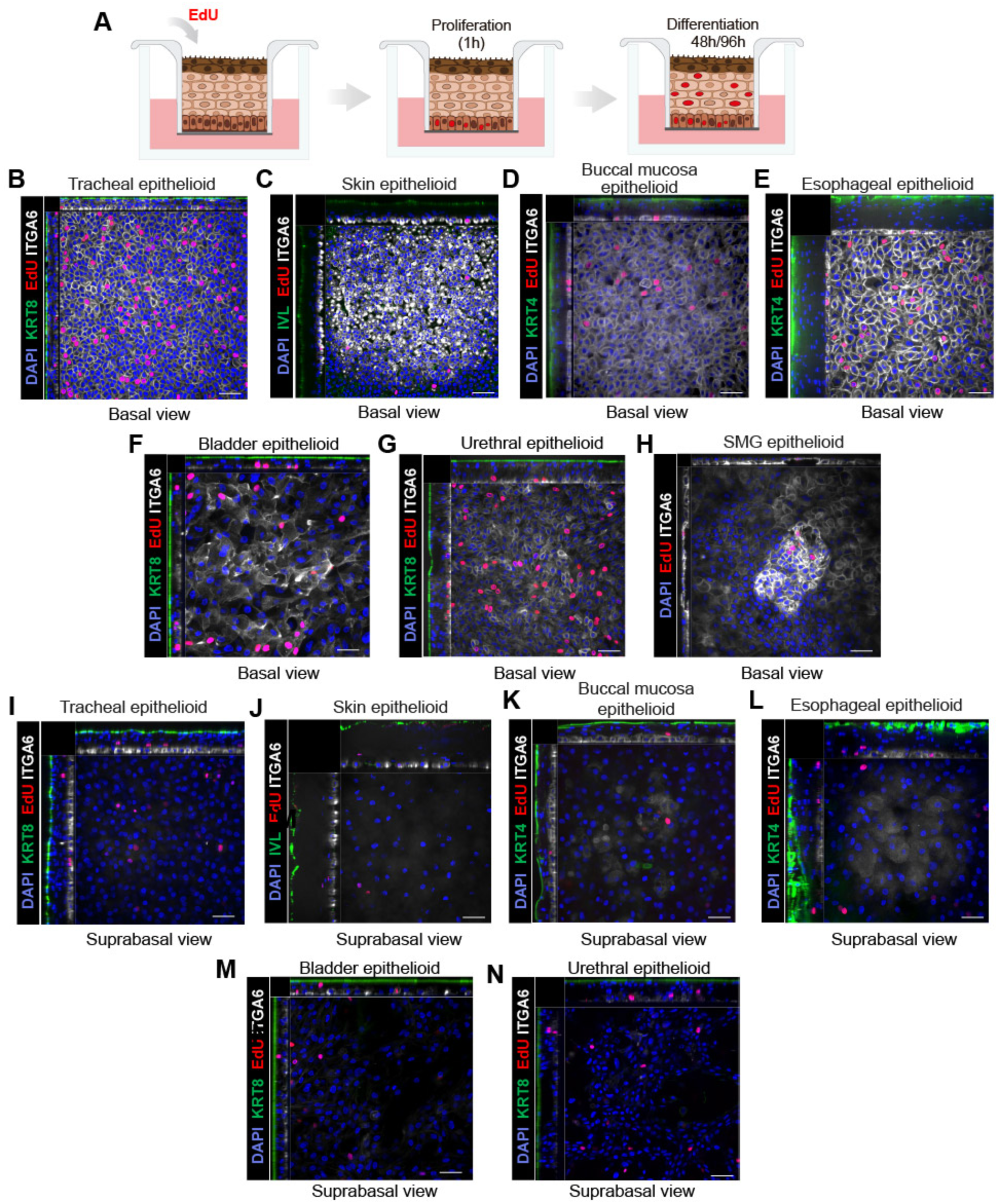
Extended data, Human epithelioids recapitulate native tissue dynamics of cell proliferation and differentiation. **(A)** Schematic of the EdU labeling assay used to measure cell proliferation (1-hour post-EdU incorporation) and differentiation (48–96 hours post-incorporation). **(B – I)** Confocal images showing basal planes of human tracheal **(B)**, skin **(C)**, buccal mucosa **(D)**, esophageal **(E)**, bladder **(F)**, urethral **(G)**, and submandibular gland **(H)** epithelioids. EdU+ cells are shown in red. Apical and basal compartments are labeled for keratins (green) and ITGA6 (white), respectively. Nuclei were counterstained using DAPI (blue). Scale bar, 50 μm. **(J – O)** Confocal images showing suprabasal planes of human tracheal **(J)**, buccal mucosa **(L)** and esophageal **(M)** epithelioids 48 hours after post-EdU incorporation, and skin **(K)**, bladder **(N)** and urethral **(O)** after 96 hours. EdU+ cells are shown in red. Apical and basal compartments are labeled for keratins (green) and ITGA6 (white), respectively. Nuclei were counterstained using DAPI (blue). Scale bar, 50 μm.

**Figure S6:**
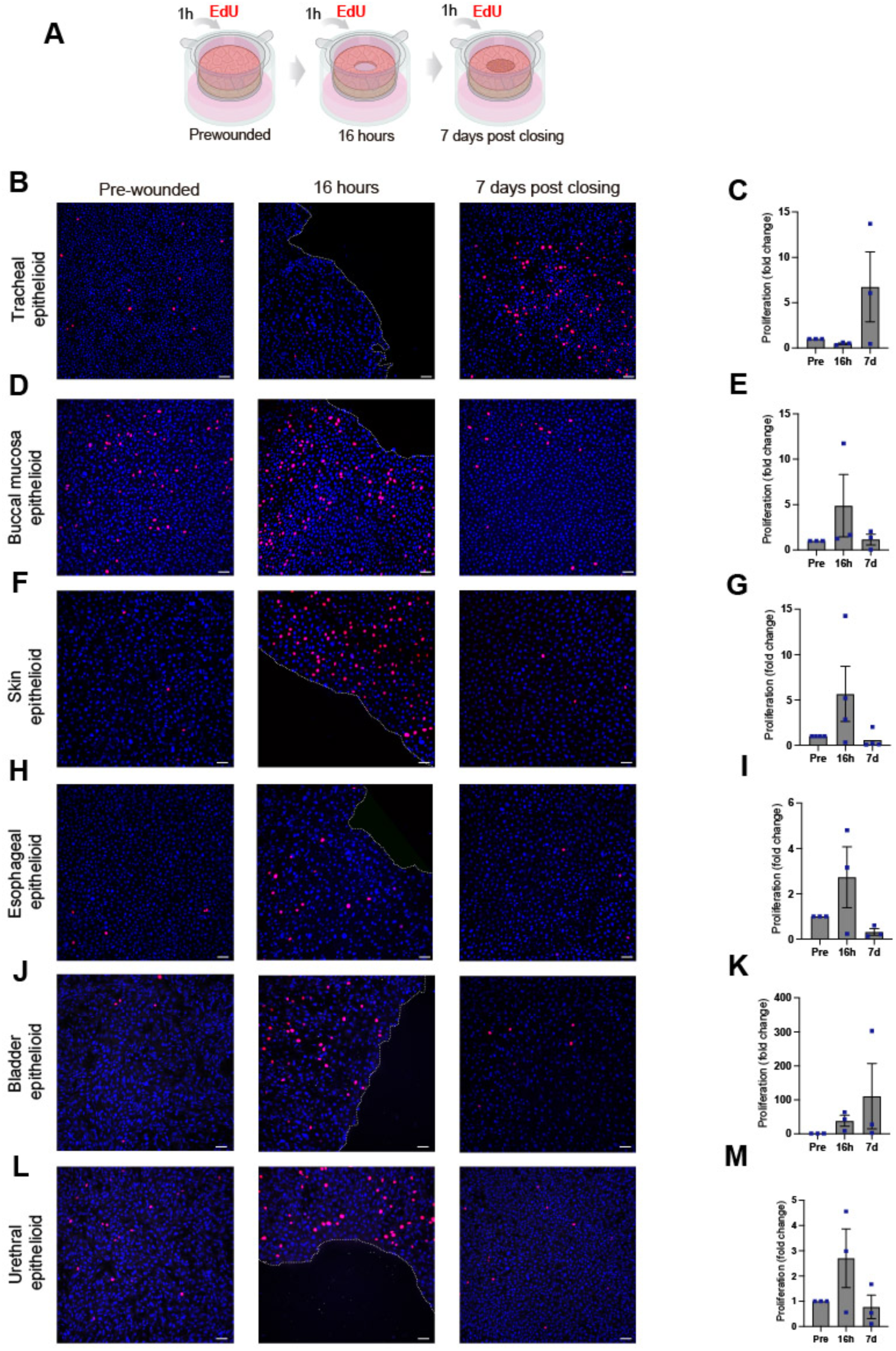
Extended data, Human epithelioids maintain barrier homeostasis and wound healing capacity after injury. **(A)** Schematic of the EdU labeling assay used to assess proliferation in human epithelioids during the wound healing process. **(B, D, F, H, J, L)** Representative confocal images showing basal views of human tracheal **(B)**, buccal mucosa **(D)**, skin **(F)**, esophageal **(H)**, bladder **(J)** and urethral **(L)** epithelioids at baseline, 16 h post-wounding, and 7 days after wound closure. EdU^+^ cells are shown in red. Nuclei were counterstained with DAPI. Scale bar: 50 μm. **(C, E, G, I, K, M)** Total number of EdU^+^ cells per mm^2^ calculated in human tracheal **(C)**, buccal mucosa **(E)**, skin **(G)**, esophageal **(I)**, bladder **(K)** and urethral **(M)** epithelioids at baseline, 16 h post-wounding, and 7 days after wound closure. Results are expressed as mean plus SD from at least three different donors.

**Figure S7:**
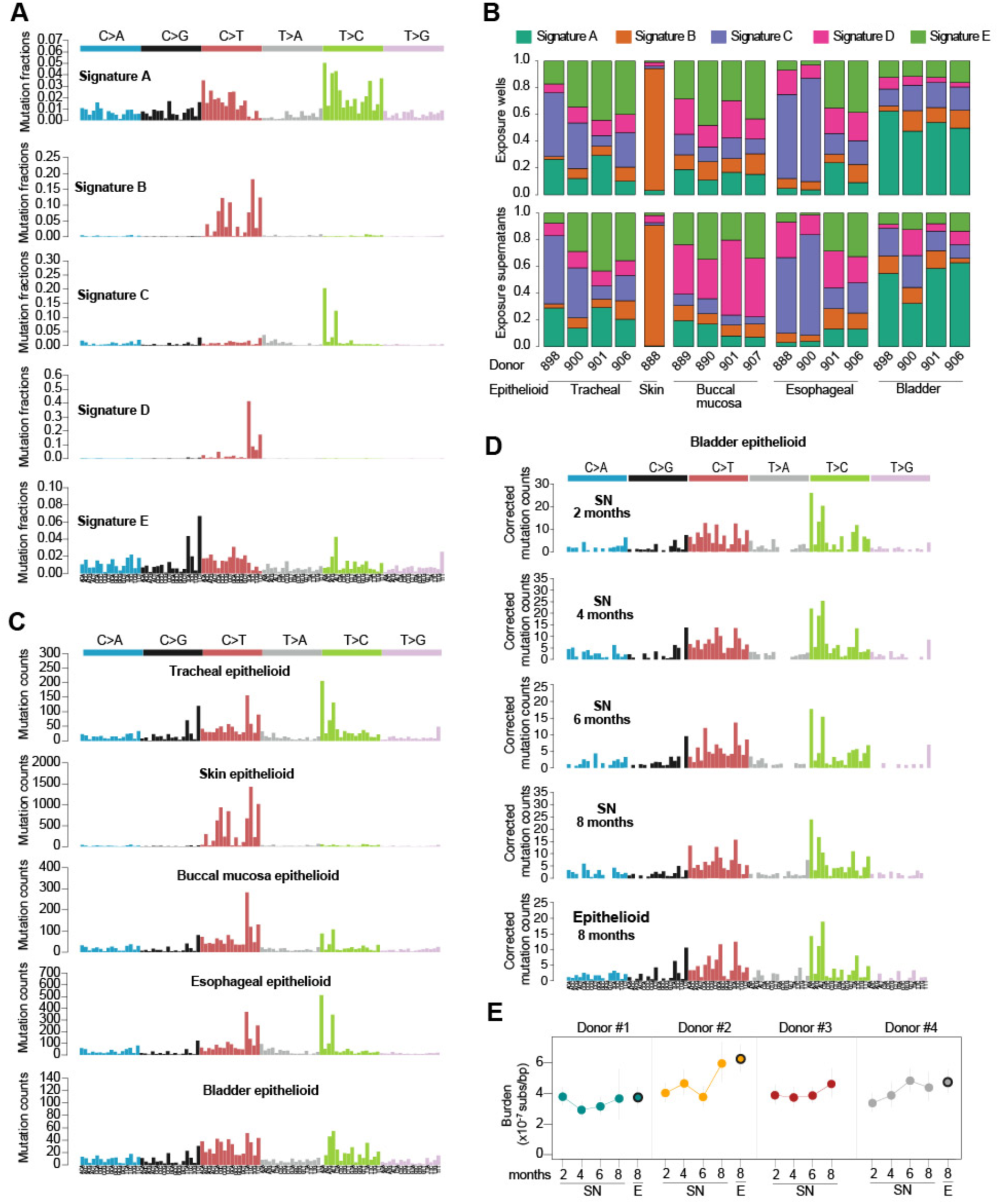
Mutation rates and signatures of human epithelioids. **(A)** Five de novo extracted mutational signatures from human epithelioids. **(B)** Signature contribution in bulk cultures (wells, top) and supernatants (bottom) of human epithelioids. **(C)** Mutational spectrum of each human epithelioid culture (only wells shown). **(D)** Mutational spectrum evolution in human bladder epithelioid supernatants and comparison with their patient matched 8-month-old epithelioid bulk cultures. **(E)** Substitution burden estimates in supernatants (SN) across 8 months in four biological replicates and comparison with their patient matched 8-month-old epithelioid bulk cultures (E, black circles).

## Resource availability

### Lead contact

Further information and requests should be directed to David Fernandez-Antoran

### Materials availability

This study did not generate new reagents.

### Data and code availability

The code and data files underlying the results of the study are available from the GitHub code repositories https://github.com/fa8sanger/Epithelioids/ and https://github.com/ishida-md/10x_Epithelioid.

Sequencing data will be deposited at EGA.

## Acknowledgments

We are grateful to the families of the deceased organ donors and to the patients with head and neck cancer for their generous consent to the use of human tissue for this research. We also thank the Cambridge Biorepository for Translational Medicine for providing access to these samples.

We thank all staff and members of the Gurdon Institute, in particular the Imaging facility for their support with confocal microscopy and image analysis, and the media preparation and administrative teams for their continued assistance. We also thank Prof. Charlotte Coles, Director of RadNet Cambridge, and Ms. Nicola Le Blond, Project Manager of RadNet Cambridge, for their continuous support and help.

This work was supported by Cancer Research UK RadNet grant n° C17918/A28870, Isaac Newton Trust / Wellcome Trust / ISSF / University of Cambridge Joint Research Grant, joint National Centre for the 3Rs - Cancer Research UK award n° NC/X000885/1, Wellcome Trust Core funding grant 203144/A/16/Z at the Gurdon Institute, University of Cambridge and the Spanish Ministry of Science and Innovation and the State Research Agency (AEI) under grant PID2024-157120OB-I00 at the Aragon Health Research Institute, to D.F-A and Instituto de Salud Carlos III through the Fondo de Investigación en Salud Projects “PI21/00441” and “PI24/00969” and co-funded by European Union (ERDF, “A way to make Europe”), to A.J.S.

I.M., F.A., A.J.R.L., Y.I., M.P. are funded by the Wellcome Trust, the Dr Josef Steiner Cancer Research Foundation, and Cancer Grand Challenges. P.A.N. was funded by a Cambridge NIHR BRC Capacity Building award (RHZB/307; grant no. G122835).

CUH Human Research Tissue Bank is supported by the NIHR Cambridge Biomedical Research Centre (grant no. NIHR203312)

## Author contributions

Conceptualization: DFA and IM. Experimental design: DFA, IM, ISF, APD and JAVL Investigation: ISF, APD and JAVL performed all experimental work and data acquisition. FA analyzed tNanoseq data. YI analyzed scRNA-seq data. MB performed sample processing, image analysis for TEM and SEM and statistical analysis, with technical support from MSN. MP performed preliminary scRNA-seq analysis. AL, PN and AO contributed to DNA processing and preliminary analysis. RB developed new software. MM and KM contributed to initial culture optimizations. CSS performed scRNA-seq. Resources: CBM, MC, AB, GH, WI and GB provided salivary gland tissue. JAT performed pathology analysis. KTM provided all human samples used across the study. Supervision: RVT supervised CSS. RJ supervised GB, CBM, MC, AB and WI. KSP and JM supervised KTM. AJS supervised MB. IM supervised FA, YI, AO, MP, AL and PN. DFA supervised ISF, APD, JAVL, MM, KM and RB. Writing: DFA and IM wrote the manuscript, with contributions from ISF, APD, JAVL, FA, YI, MB, MP and AJS. Review and editing: all authors reviewed and edited the manuscript.

